# Training a neural network to learn other dimensionality reduction removes data size restrictions in bioinformatics and provides a new route to exploring data representations

**DOI:** 10.1101/2020.09.03.269555

**Authors:** Alex Dexter, Spencer A. Thomas, Rory T. Steven, Kenneth N. Robinson, Adam J. Taylor, Efstathios Elia, Chelsea Nikula, Andrew D. Campbell, Yulia Panina, Arafath K. Najumudeen, Teresa Murta, Bin Yan, Piotr Grabowski, Gregory Hamm, John Swales, Ian S. Gilmore, Mariia O. Yuneva, Richard J.A. Goodwin, Simon Barry, Owen J. Sansom, Zoltan Takats, Josephine Bunch

**Affiliations:** National Physical Laboratory, Teddington, TW11 0LW; Beatson Cancer Research UK Institute, Glasgow, G12 0YN; Francis Crick Institute, London, NW1 1AT; Clinical Pharmacology and Safety Sciences, BioPharmaceuticals R&D, AstraZeneca, Cambridge, CB4 0GQ; Imperial College London, London, SW7 2BX

## Abstract

High dimensionality omics and hyperspectral imaging datasets present difficult challenges for feature extraction and data mining due to huge numbers of features that cannot be simultaneously examined. The sample numbers and variables of these methods are constantly growing as new technologies are developed, and computational analysis needs to evolve to keep up with growing demand. Current state of the art algorithms can handle some routine datasets but struggle when datasets grow above a certain size. We present a training deep learning via neural networks on non-linear dimensionality reduction, in particular t-distributed stochastic neighbour embedding (t-SNE), to overcome prior limitations of these methods.

**One Sentence Summary:** Analysis of prohibitively large datasets by combining deep learning via neural networks with non-linear dimensionality reduction.

## Main Text

When describing big data, two types of big data challenges are often discussed; data with large sample numbers such as stock exchange, marketing, and social media data, and data that contains a large variable space (high dimensionality), such as genomics, transcriptomics, and mass spectrometry based omics (proteomics and metabolomics).

Some methods have high sample numbers but low numbers of variables (such as electron microscopy), and as such analysis is limited to review of single or composite images. On the other hand, there are methods that produce very high dimensional data, but the sample numbers are sufficiently low that memory efficient analysis algorithms need not be developed. Examples of these include many standard omics data such as genomics^1^, metabolomics^2^, proteomics^3,4^ and transcriptomics^5^. Then there are methods that combine both of these challenges having vast numbers of variables from large sample numbers. These types of data present the greatest challenges for analysis and information extraction. Examples of these techniques include spatial and single cell transcriptomics^6^, hyperspectral imaging^7^, chromosome conformation capture (Hi-C)^8^, mass spectrometry imaging (MSI)^9^, and imaging mass cytometry^10^. Common tasks required to analyse these data include image segmentation, dimensionality reduction, and classification as a means for interpretation, and visualisation^11-13^.

However, when datasets are particularly large, many of these algorithms fail due to lack of memory (RAM), and in extreme cases even loading the full dataset into memory becomes prohibitive. Examples of these include large Hi-C data, hyperspectral imaging, and mass spectrometry imaging (MSI)(Figure 1).

**Figure 1.**
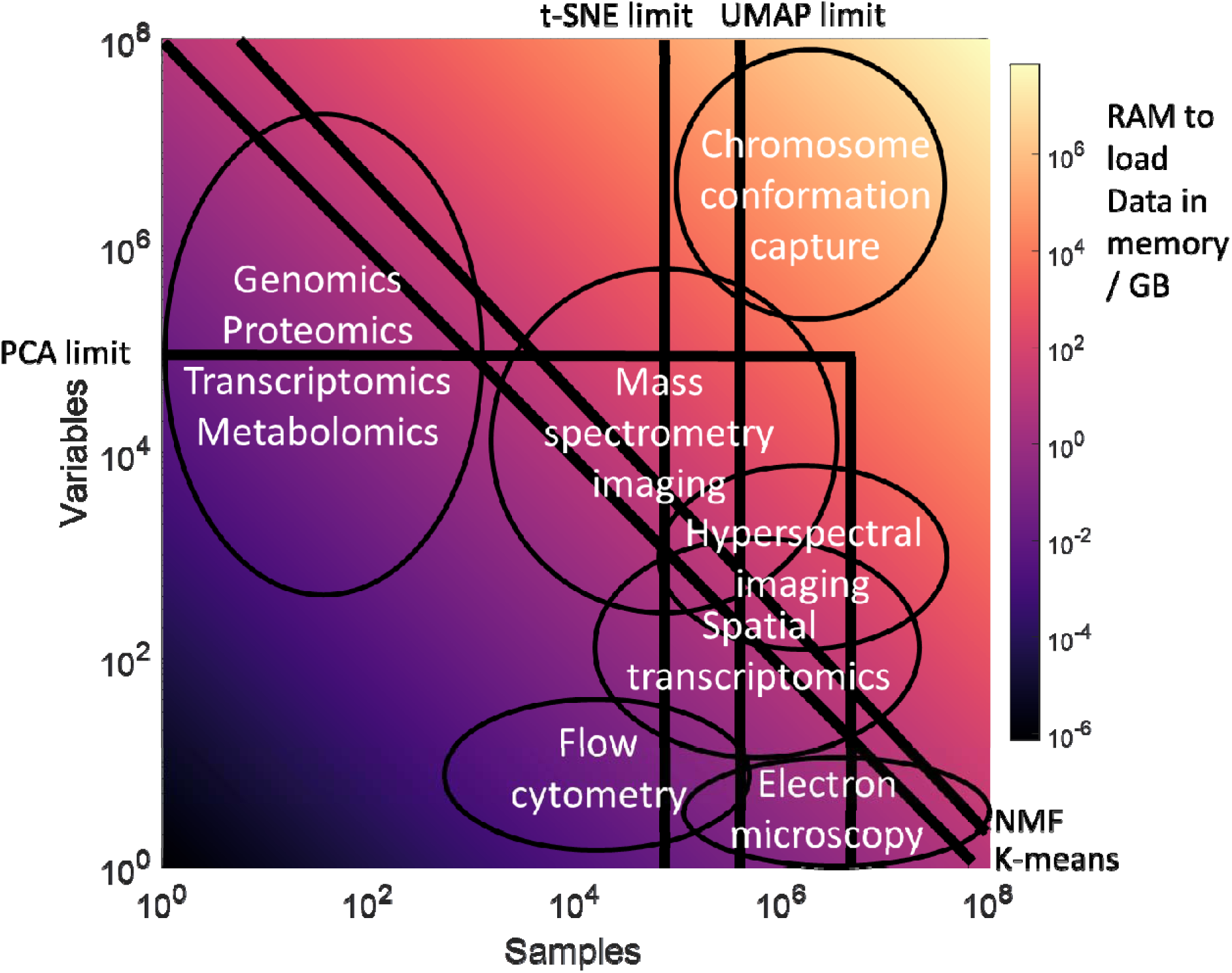
The range of sample numbers and variables associated with common and emerging tools in biomedicine. Colour scale indicates the amount of RAM required to load data of the corresponding size into memory, and lines indicate the computational limit for principal component analysis (100,000 variables), t-SNE (50,000 samples) and UMAP (500,000 samples).

Non-linear dimensionality reduction via methods such as t-SNE and uniform manifold approximation and projection (UMAP) provide insightful feature extraction, and are commonly used in analysis of omics data^7,14-16^, and other areas where sample numbers remain relatively low. These algorithms have been shown to outperform other linear dimensionality reduction methods in a number of different areas including document clustering^17^, genomics^16,18^ transcriptomics^6^, and visualisation tasks^14,19^.

There are three major limitations of both of these algorithms however:

- Applying t-SNE and UMAP to large sample numbers requires a lot of computational resources. This is because the memory usage (space complexity) and calculation time (time complexity) scales by the number of pixels (*n*) squared for t-SNE^20^, or by *n*^*1*.*14*^ for UMAP^21^. This means it is not feasible to perform reduction on large datasets using standard computers in a reasonable timeframe.
- The second fault with these methods is that typically the whole dataset is required in order to create the low dimensional representation. This means that in order to reduce additional data alongside existing data, a new instance of reduction must be performed using a combined dataset of existing and additional data. This limitation, combined with the high complexity, both in time and memory, mean that it is not practical to repeat these methods often.
- Unlike algorithms such as principal component analysis (PCA) and non-negative matrix factorisation (NMF)^22^, there is no way to determine the high dimensional contributions towards the low dimensional representation. It is critical in some areas to not only extract features, but also to relate these features to the high dimensional information.

The issue of computational complexity has been addressed in t-SNE by several different approaches. The parametric t-SNE algorithm has similarities with the methodology we are proposing in that it uses a neural network based training using the t-SNE cost function^23^. However unlike our proposed method this method does not perform t-SNE and the reduction itself is performed by neural networks. Other tree-based and fast Fourier Transform approximation methods to the pairwise distances of pixels have been investigated^24^. These are able to reduce both the memory requirement and computational complexity to O(*n* log *n*) depending on the depth of the tree. Unfortunately this is still insufficient for extremely large datasets^25^.

Abdelmoula *et al*. recently demonstrated the use of a hierarchical stochastic neighbour embedding method using landmark features to embed and visualise large 3D mass spectrometry imaging datasets^26^. This allowed t-SNE dimensionality reduction to be performed on datasets with over 1 million pixels such as 3D MSI data. Along with this, Boytsov *et al*. recently proposed the use of local interpolation with outlier control t-SNE (LION-tSNE) to embed newly acquired data into previously t-SNE mapped space^27,28^. As well as allowing t-SNE to be performed on large datasets, this also allows new data to be incorporated into the embedded 2D or 3D space. Although this method demonstrates the power of mapping data into a previously determined lower dimensional space, outlier data is randomly assigned to new locations, and so there is no way to understand how outlier data are related to the existing data.

Subsampling of data has been used to improve algorithm efficiency in many different fields for tasks such as clustering and PCA^9,29^, as well as t-SNE in the viSNE algorithm^30^. By subsampling the data and applying dimensionality reduction or clustering to a subset of it, more efficient data analysis can be performed. The limitation with this for t-SNE and UMAP is that there is no mathematical transformation that can then be applied to the remaining data to embed it into the low dimensional space.

Deep learning via neural networks can be used to learn mathematical transformations to go from high to low dimensional space. These can be either supervised such as those used in classification problems^31,32^, or unsupervised, such as stacked and variational autoencoders^33,34^. Neural networks are often applied in classification problems, demonstrating greater accuracy than linear approaches ^35^.

Recently, there has been a development of methods using neural networks and autoencoders to perform dimensionality reduction itself. In particular, methods such as parametric t-SNE, VAE-SNE and net-SNE have used the Kullback-Leibler divergence used in t-SNE as an optimisation function for autoencoders to produce similar results to t-SNE^36,37^. Other methods such as IVIS use Siamese neural networks with triplet loss function to preserve local and global similarity in the reduced space^38^. Recently Abdelmoula *et al*. demonstrated the usefulness of these types of approaches to return spectral information from MSI data as well their capability to handle large datasets^39^.

To date, the majority of developments in neural network based non-linear reduction have been aimed at performing the non-linear reduction optimisation using cost functions derived from existing methods such as t-SNE. Espadoto *et al*. showed that neural networks can be trained on t-SNE and UMAP projections directly, and demonstrated this on relatively small testing datasets^40^. Here we demonstrate the use of deep learning on subsampled non-linear dimensionality reduction using t-SNE and UMAP to extract features from large complex biological datasets. We deploy this method on datasets from different fields including Hi-C, hyperspectral imaging, and MSI.

## Results

### Description of the method

Our approach is to take a small subset of the data and perform dimensionality reduction on it. The resulting dimensionally reduced data is then used as the training set for deep learning using neural networks (Figure 2). The remaining data not used for training is then embedded into the same reduced space along with any other new data subsequently measured. A similar set of networks are also trained on the reverse from the lower into higher dimensional space. This allows any hypothetical point in the low dimensional space to be returned into a predicted high dimensional data point. This algorithm includes several steps, each of which could be independently optimised, and many will not be independent of one another. A fully exhaustive comparison of these parameters is not feasible in the current work, but we will discuss the main parameters that are unique to this method. These include subset size used, method of subsampling the data, and the algorithm used when training the neural networks themselves. In all other cases, optimal parameters taken from prior literature have been used.

**Figure 2.**
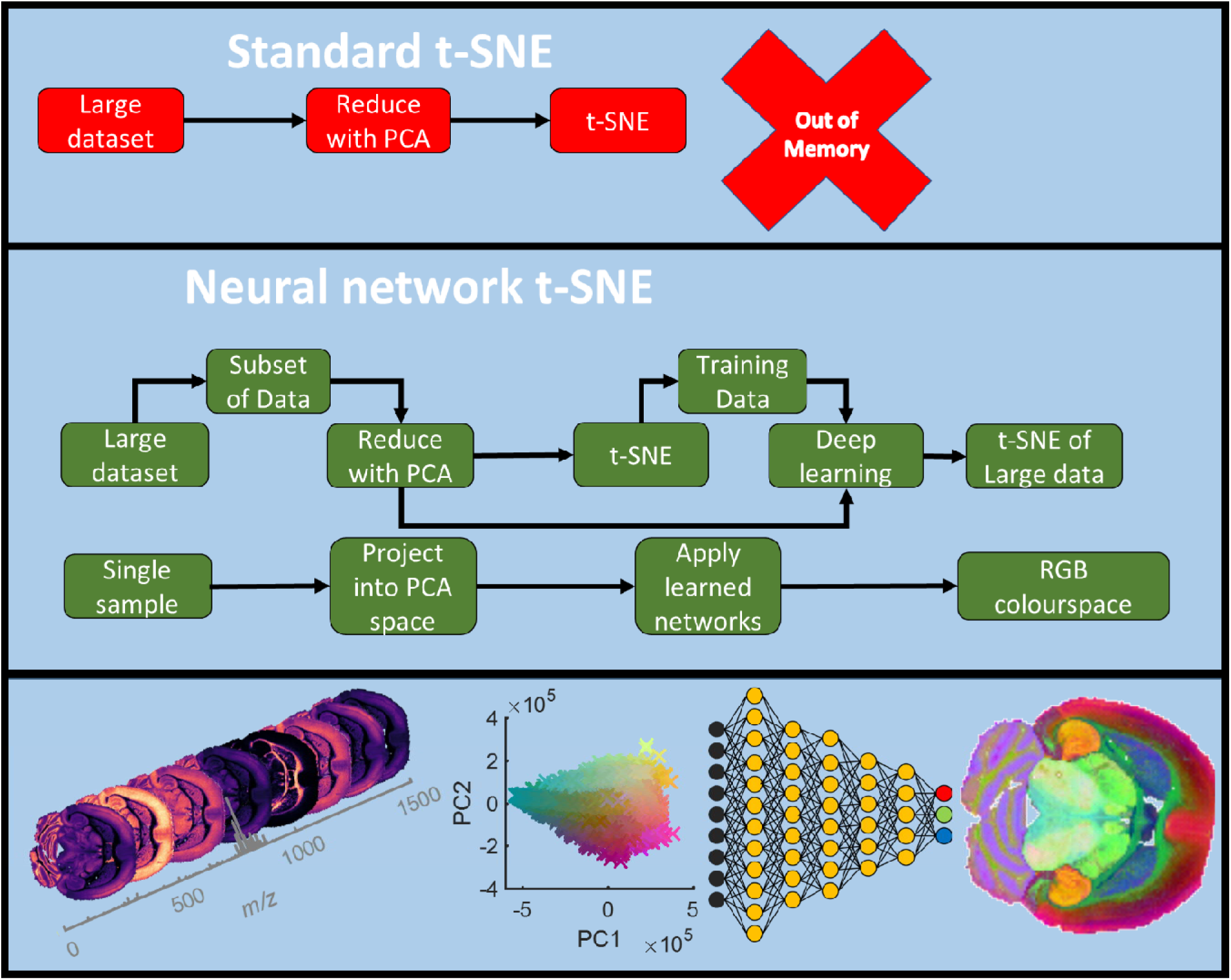
Outline of the novel workflow using t-SNE as the base dimensionality reduction method. A large dataset that cannot have t-SNE performed on it is subsampled, and t-SNE is performed on the subset of data. The resulting embedding is then used as training data to train deep learning via neural networks to go from the high to low dimensional space. The networks can then be applied to remaining data (and any new data) resulting in reduction of 5 the full dataset into an approximation of t-SNE.

### Example case studies

#### Large data t-SNE

The foremost challenge is to apply t-SNE to larger datasets, broadening the applicability of this method. This has been addressed by hierarchical t-SNE and LION-tSNE described previously^26,27^. Using NN t-SNE is another way to achieve this with greater efficiency than these existing methods. We demonstrated this using a publicly available chromosome conformation capture dataset containing over 2 million samples^8^. This method is able to rapidly produce informative results on these large data that could not be analysed using standard t-SNE methods (Figures 3 and S1). Hi-C is a DNA sequencing-based method for probing the 3D structure and interactions of chromosomes. This type of data is notoriously challenging to load into RAM and process, even on modern computers. We performed a three-dimensional embedding of the entire Hi-C dataset for a human GM12878 cell line^8^ using the highest possible resolution of 1kb. The resulting embedding captured the spatial relationships between the chromosomes encoded in the Hi-C dataset (Figure 2a). Not surprisingly, the 22 chromosomes exhibit more intrachromosomal interactions than interchromosomal interactions. Moreover, the approach captured more hierarchical nature of the human nucleus, such as division into discrete A/B sub-compartments, described previously^8^. These sub-compartments are regions of chromosomes which harbour different epigenetic marks and display a tendency to physically interact with regions in the same sub-compartment type. After annotating our 3D embedding with the H3K27ac ChIP-seq signal (H3K27ac is an epigenetic mark linked to active gene expression), we observed that regions of high transcriptional activity co-cluster together in the 3D space (Figure S1). This highlights the hierarchy and complexity encoded in our unsupervised embedding and suggests that using this approach can further aid studies of the human genome architecture, for example in identifying novel genome sub-compartment types which could be missed using lower resolution data. In order to perform t-SNE on these data, over 80 TB of RAM would be required, making these analyses infeasible on these types of datasets. Using the NN t-SNE method, less than 10 GB are required at any one time to segment these data.

**Figure 3.**
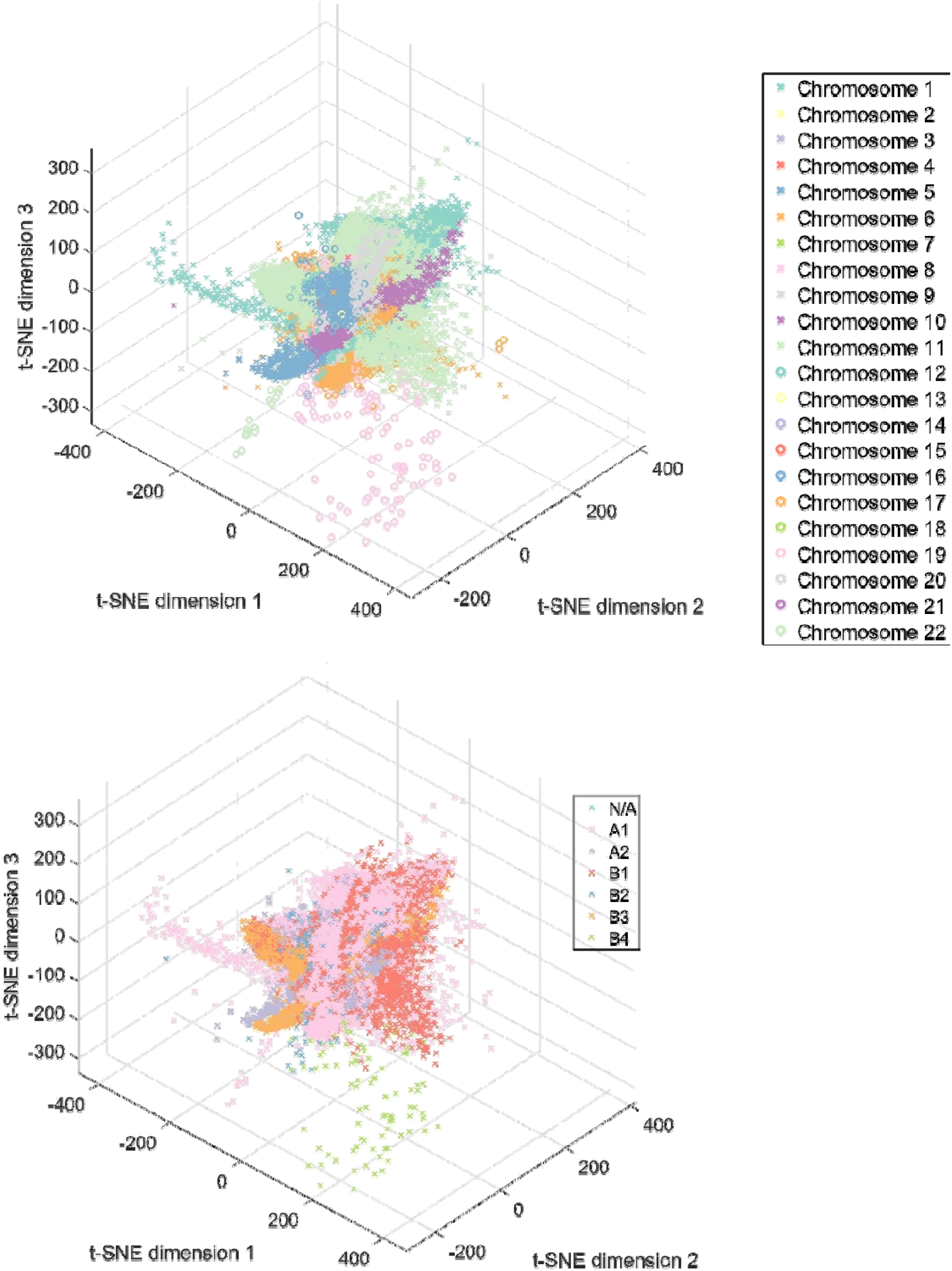
Results of 3D embedding from NN t-SNE dimensionality reduction on Hi-C data containing over 2 million samples labelled by chromosome (A), or interactions (B). Using standard t-SNE approaches would not be possible for either this dataset due to memory constraints, and simply loading these data into RAM is often not possible.

#### Addition of new data

While performing t-SNE on large datasets is useful, any requirement to incorporate new data would necessitate rerunning the whole process to obtain segmentation of all data together. This would likely be time consuming, produce a different colour mapping (due to the stochastic nature of t-SNE), and may exceed the memory constraints of this algorithm. Using the NN t-SNE approach, new data can be embedded into the low dimensional space of the previously segmented data. This can either be used to find data similar to the existing data, or data that is different from the training data for outlier detection.

To demonstrate segmentation of new data into the same RGB colour mapping, two datasets were selected; a hyperspectral imaging data from a series of faces from the isetbio database (http://52.32.77.154/repository/isetbio/resources/scenes/hyperspectral/stanford_database/faces3m/.accessed30/08/2019) containing over 30 million (1m per image) samples and 160 variables, and MALDI MSI images of 30 coronal serial sections of drug treated mouse brain glioblastoma model were taken (500,000 samples, 4,000 variables).

To analyse the hyperspectral imaging data, a subset of the data from the face of one person was selected for training, and the learned networks were applied to the images from other datasets. The pixels from the same features were easily and quickly segmented and discerned as similar to one another demonstrating that this method can be applied to many datasets sequentially (Figure 4). The rapid and effective evaluation of features in hyperspectral imaging is important for areas such as biomedicine where it may be used for classification purposes^41^, or in satellite imaging where algorithms will need to match ever increasing data rates afforded by new instrumentation^42^.

**Figure 4.**
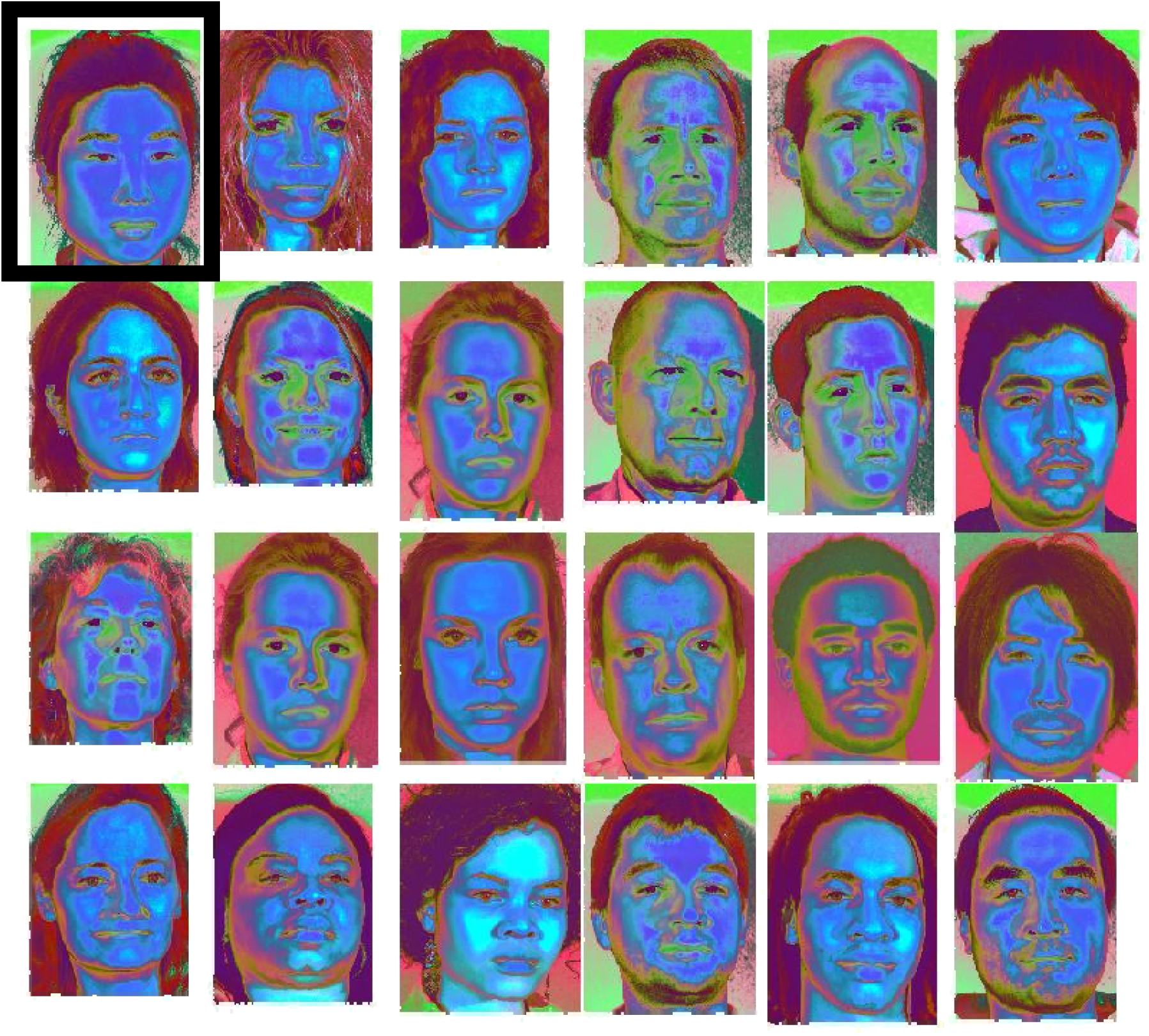
Visualisation of the results from NN t-SNE trained on a single hyperspectral image (highlighted by black box), and then applied to the data from the remaining images showing a consistent segmentation of hyperspectral facial features in a unified colour scheme. We can see clearly that pixels from the same features such as face, hair and background are consistently 5 segmented, allowing for rapid extraction of common features between datasets.

In the MSI images, data from one of the tissue sections (containing the tumour) were taken as the training data, t-SNE was performed on it, and neural networks used to learn a transformation from the original to t-SNE space. The trained network was then applied to the data from the remaining 29 serial sections, and a consistent segmentation of the same anatomical features into the same colours can be seen (Figure 5a). As well as producing consistent segmentation of anatomy, the application of the network to the new data took around 1.5s per dataset, far faster than the data acquisition itself. The consistent segmentation provides an excellent platform for registration and resulting 3D segmented image generation (Figure 5b).

**Figure 5.**
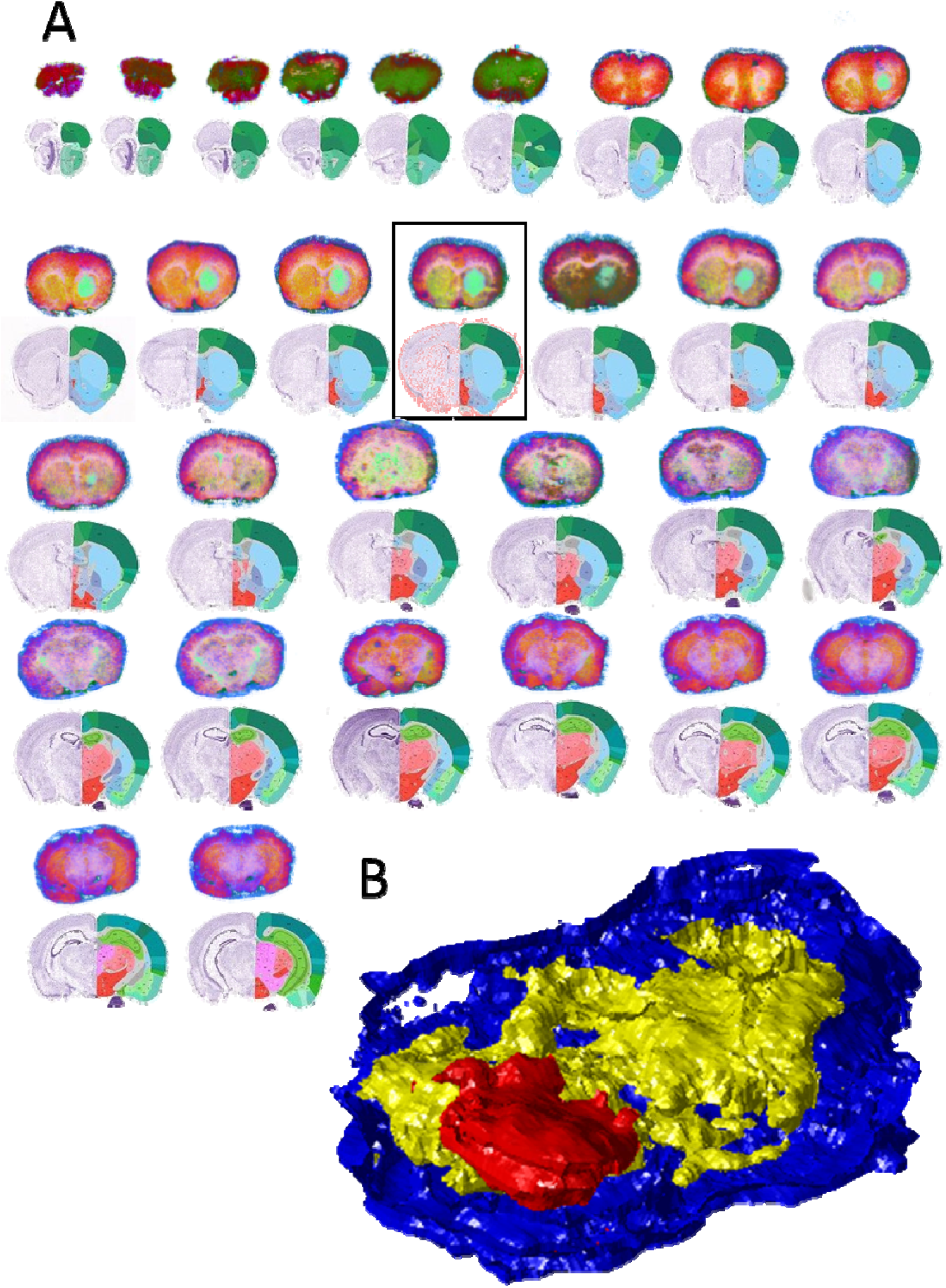
Results of segmentation using neural network t-SNE trained on MSI data from a single section of murine brain (highlighted in red) containing glioblastoma (segmented in light green). The data from the remaining sections were then reduced using the trained networks to segment the same anatomies in the same colour scheme. By embedding data into the same colour space similar features can easily be identified, and the results of this can easily be used to perform registration to generate 3D representations such as shown in (B).

In addition to consistently segmenting the same anatomies of similar data, new data that is dissimilar to all existing data can be identified. This is demonstrated on the same MSI data from the serial sections of mouse brain. By training the network on t-SNE applied to a section without tumour present, the tumour tissue and the cerebellum region is still identified as being different from all tissue within the training data (Figure S2). This means that this approach can be used to identify outlier data from the training data, could be used as a quality control metric, or be used to identify data of interest for additional analysis.

The consistent embedding provided by NN t-SNE can also be applied to non-imaging mass spectrometric data. A promising new area of mass spectrometric research is rapid evaporative ionisation mass spectrometry (REIMS)^43,44^. This has the potential to classify tissues during surgeries in almost real time. In addition to this, it can be used for rapid cell profiling^45^. NN t-SNE analysis was performed on REIMS data acquired from cell pellets extracted from the small intestine and colon of mice with different oncogenetically relevant mutations. There is a much clearer separation in the genetic types observed by this method as compared to PCA (Figure S3). These data were then used as a basis for classification by linear discriminant analysis (LDA). The classification based on the NN t-SNE reduced data shows much higher accuracies as compared to data reduced by PCA (Rand index 0.93 compared to 0.83, Table S1), indicating that this may be a better dimensionality reduction method to apply prior to classification. It is important to note that a normal t-SNE approach would be inadequate for classification because new data cannot be projected into the reduced space. The clinical application of this approach is clear - it allows real-time stratification of patients/mice into groups based upon multidimensional datasets, rather than binary segregation based upon presence or absence of individual dominant oncogenic mutations. The latter approach has broadly failed as a both as stratification strategy and has little prognostic value, and in the case of colorectal cancer, has been superseded by more advanced molecular subtyping approaches (genomic, transcriptomic), which while inherently more representative of the underlying biology of a tumour, are both costly and time-consuming.

#### Returning spectral information using MSI data

One of the biggest limitations of t-SNE as compared to other dimensionality reduction methods like PCA and non-negative matrix factorisation is that no spectral contributions driving the differences between features can be obtained. By reversing the neural networks, we can obtain a transformation to go from any three-dimensional RGB value (or any other t-SNE reduced space) into a spectrum. By performing this for red ([1 0 0]) green ([0 1 0]) and blue ([0 0 1]) we can obtain the relative spectral contributions along these dimensions to our image segmentation. Alternatively, we can segment the reduced space and calculate centroids in the reduced space to determine the spectrum that would be derived from that region. We demonstrate this capability using DESI MSI data of mouse mammary gland tumours containing 90,000 samples and 2,000 variables.

The oncogene MYC is amplified in approximately 15% of breast cancers and is associated with poor prognosis. Oncogenes such as MYC define metabolic vulnerabilities in tumours. ErbB2/HER2 is overexpressed in 20% percent of human breast cancers and correlates with tumour chemo-resistance and poor prognosis^46^. Mammary gland tumours were generated in transgenic mouse with ectopic expression of either MYC or ErbB2 under the control of mouse mammary tumour virus (MMTV) promoter^47,48^. Resulting tumours were harvested at max 1.2 cm. These two oncogenes result in distinct metabolic profiles. Normal mammary glands (NMG) from healthy mice were also analysed. For each phenotype two biological replicates were analysed in a single acquisition.

NN t-SNE of these data following background subtraction (Figure 6) clearly segments the three samples (MYC and ErbB2 tumours and NMG) in low dimensional space (Figure 6a-b). Bounding boxes describe distinct areas including leached fatty regions from the NMG, and the majority of pixels in the MYC or ErbB2 tumours (Figure 6c).

**Figure 6.**
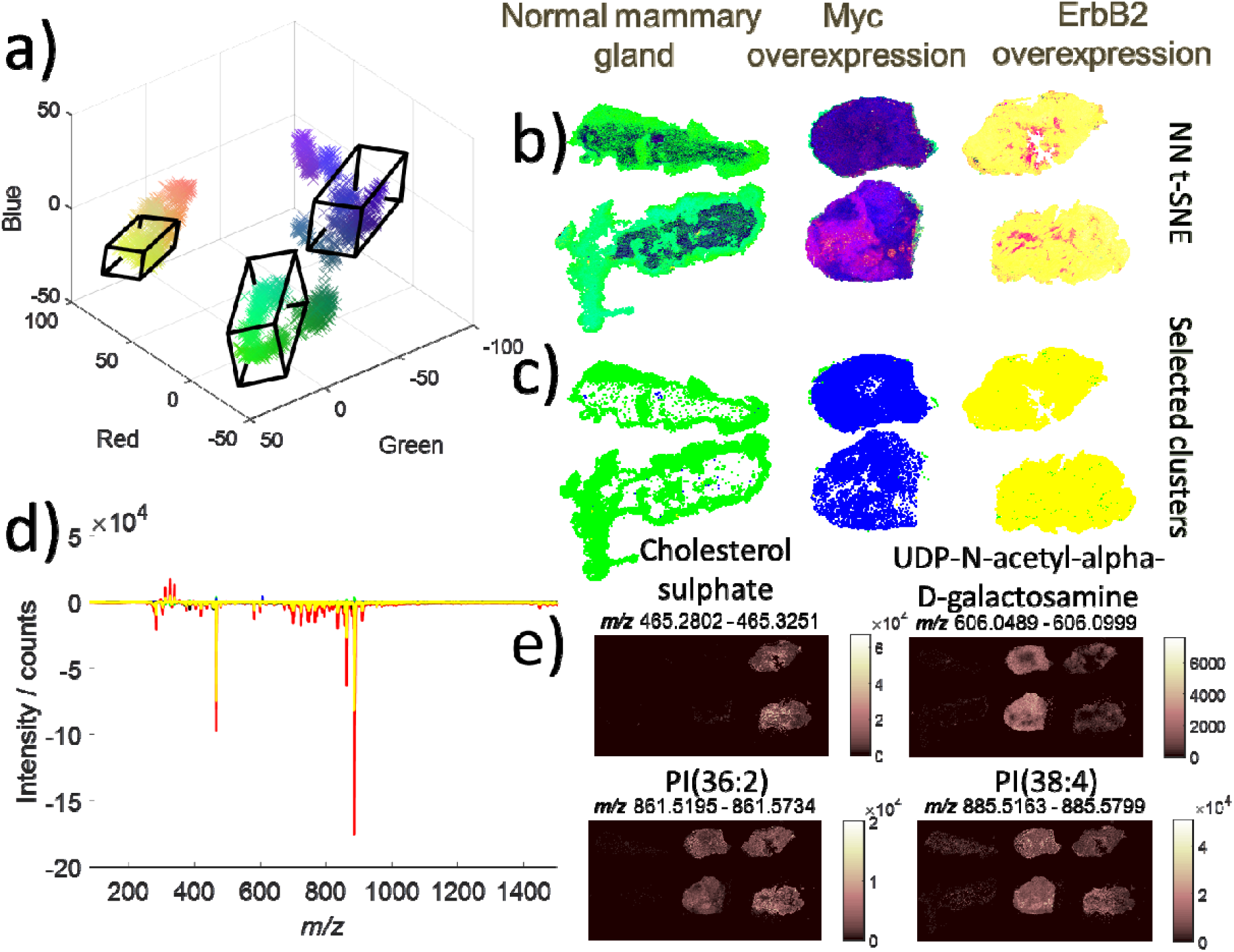
Results of NN t-SNE applied to MSI data from genetically engineered breast cancer models. The three genetic models are clearly differentiated in both the scatter plot (a) and image (b). These can be segmented to give masks for the different models (c) and the underlying spectral drivers can be identified (d) and their corresponding ion images generated (e).

While such spatial segmentation is useful, in order to provide biological interpretation we must interrogate which spectral features are driving the separation in low dimensional space. We generated the spectral representation for selected corners of the low dimension space (red, green, blue and yellow) (Figure 6d). Spectral representations may include positive and negative values indicating their relative contribution at the selected coordinates. Selected peaks with strong contributions to the spectra representation for each selected low dimensional coordinate were matched by exact mass to HMDB-v4.0 database^49^ and putative annotations made (Figure 6e). Cholesterol sulfate, a structural component of cellular membranes, has been previously identified in DESI MS experiments as a potential biomarker of prostate cancer^50^. Here it is a strong contributor to the “yellow” corner of low dimensional space where pixels from the ErbB2 tumour are positioned. This appears to be a strong marker for ErbB2 tumours over MYC tumours and NMG and it localised throughout the tumour, not just in the blood and plasma-rich regions. Two peaks annotated as PI lipids are prominent in the spectral representation of the red and yellow side of low dimensional space. In the single ion images, they are localised in the MYC and ErbB2 tumours but absent in the NMG. As cell density is notably higher in the tumours than the sparse fat pad, this elevation of structural lipids in the tumours is expected. A notable contributor to the spectral representation of the “blue” corner of low dimension space is m/z 606, assigned as UDP-N-actyl-alpha-D-galactosamine. This nucleotide sugar has been implicated protein glycosylation^51^. It is present in both tumour types but elevated in MYC-driven tumours, with some heterogeneity within individual tumours observed.

#### Combining these attributes (large data consistent segmentation and spectral information)

When acquiring data from biological samples, technical and biological replicates are important. Even just performing these in duplicate, this means that for any single tissue type, four tissues may be analysed. Adding multiple genetic variants and tissue types further increases the amount of data for comparison. This was demonstrated on a MALDI dataset from the colon of four genetic variants of genetically engineered mouse models (wild-type, Apc defficient, Kras^G12D^ mutant, and dual Apc deficient with Kras^G12D^ mutation), analysed with duplicate biological and technical replicates (total of 24 tissue sections). This dataset contained 427,967 pixels and 4,000 spectral features. Using the NN t-SNE algorithm, we see that there is a clear cluster relating to the KRAS mutation containing mice (pink cluster Figure 7), a cluster which is primarily present in the mice with Apc deficient intestinal tissue (dark blue cluster Figure 7) as well as a cluster in all but the wild type mice (light blue cluster Figure 7). The main drivers of these differences were *m/z* 885.550 (light blue), 514.280 (dark blue), and 835.530, 861.550, and 887.566 (pink). These masses were then matched to the HMDB within 5 ppm mass error to provide tentative assignments of PI (38:4) [M-H]^-^ (*m/*z 885.550) taurocholic acid [M-H]^-^ (*m/*z 514.280), and PI (34:1), (36:2), and (38:3) [M-H]^-^ respectively(*m/*z 835.530, 861.550, 887.566). The PI species identified in this study have clear biological relevance, with PI(38:4) known to be the most abundant mammalian phosphatidyl inositol^52^. Phosphatidyl inositide species are critical cell signaling secondary messenger molecules whose relative abundance is both influenced by oncogene or tumour suppressor mutations (such as Apc loss or Kras mutation), and are critical for characteristic tumour cell phenotypes including unrestrained growth and survival signalling.

**Figure 7.**
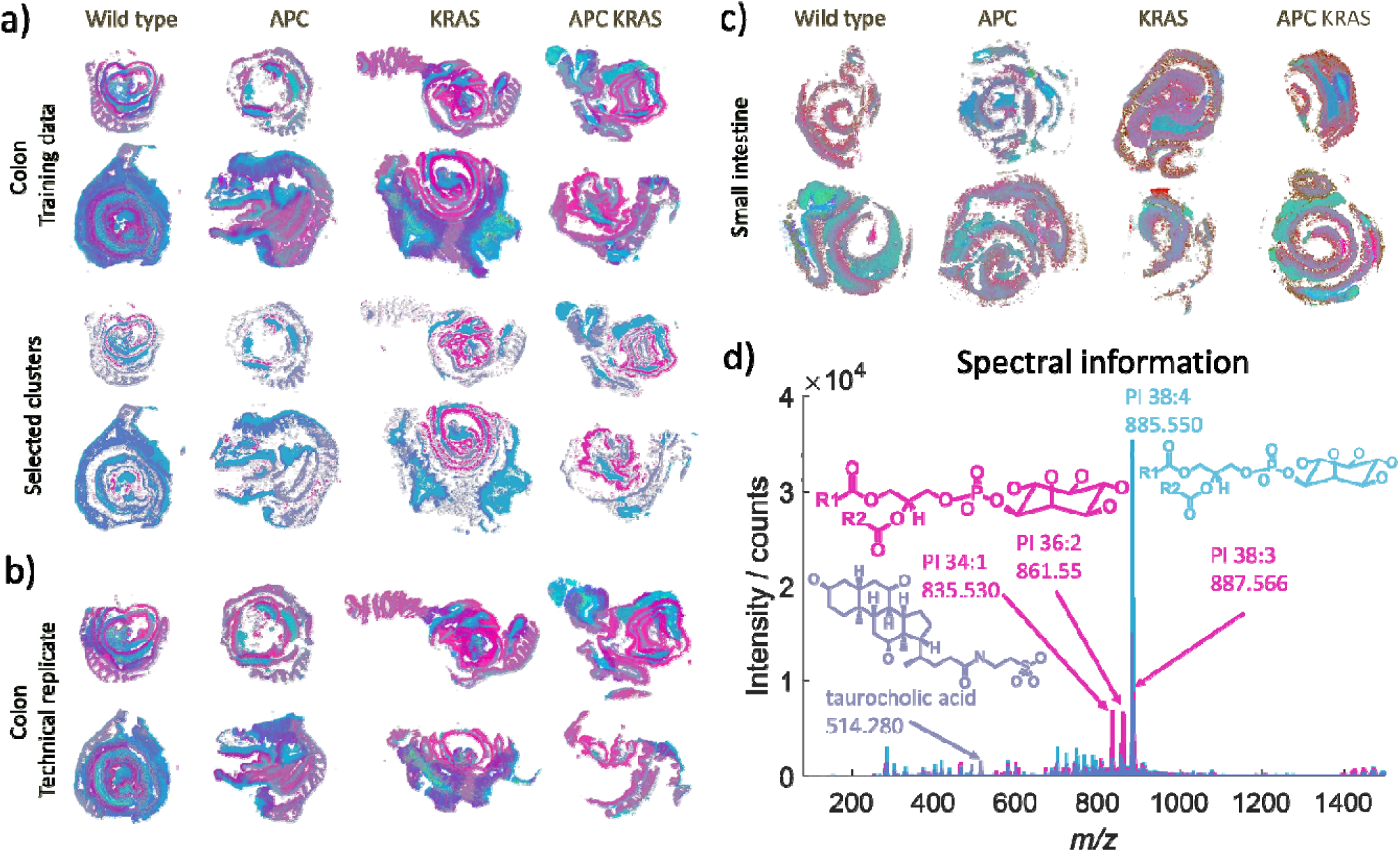
Results of neural network t-SNE applied to MSI data from biological and technical replicates from a genetically engineered mouse models for colorectal cancer. The RGB colour space from the NN t-SNE clearly differentiates the genetic differences based on difference in the metabolites detected by MSI. The pink cluster is observed primarily in the KRAS mutation containing tissues, while the purple cluster is prevalent in the APC deletion containing tissues. These clusters are driven by phosphatidyl inositol lipids and taurocholic acid respectively. This segmentation is consistent in technical replicated (b), and additional data from small intestine can be compared with comparable colour segmentation (c).

Similarly, the identification of the purple cluster associated with Apc deficiency driven by a secondary bile acid such as taurocholic acid may give valuable insight into the comparative biology of transformed versus normal intestinal enterocytes and relate to the critical role played by the gut microbiome in initiation and progression of colorectal cancer^53^.

#### Parameter optimisation

There are a several considerations to be taken into account when applying neural network t-SNE to MSI datasets. One of these is the effect of the size of the subset on the accuracy of the segmentation. Due to the stochastic nature of t-SNE, performing it on different subsets will give different results each time, making evaluation of the segmentation challenging. Therefore, a smaller dataset was taken, on which t-SNE was performed on the whole data. Subsets of the final reduced data were then taken as the input for training a neural network. The result of the neural network trained t-SNE was then compared to the t-SNE on the complete dataset by means of the correlation of the resulting reduced three-dimensional space. This was performed by subsampling the data both randomly and in an ordered manner.

The effect of random or ordered subsampling seems to have little to no effect on the end result in this instance (Figure S4 and S5), as both methods converge to give the same correlation to the original t-SNE (within error bars) from around 10% subset size, and similar anatomical segmentation. Efficient subsampling to give subsets that are representative of the whole data would be an interesting topic for a future study. For example, in image analysis, Sobol series sampling has been used to minimise the unsampled areas of the image when performing a subsampling for PCA^29^.

To analyse the effect of subset size on the whole algorithm, including the t-SNE portion, well known samples are required. Towards this end, sagittal, transverse and coronal mouse brain tissues were chosen as they have very well-defined anatomical features. These were acquired with a variety of different pixel sizes (100, 45 and 20 µm) to give a total number or pixels ranging from 10,000 to 100,000. NN t-SNE was then applied to these datasets using subsets ranging from the whole data down to 0.1% of the data (Figure S6). As might be expected, the larger the subset of data, the better the neural network training. The performance is not linear however, and below a subset size of 1,000 pixels the segmentation performs very poorly compared to the expected anatomical features (Figure S7). The measure of image autocorrelation introduced for MSI by Smets *et al*.^*21*^ was also used to evaluate the results from the subset size comparison on the sagittal brain dataset, and similarly we see a large decrease in the correlation between subsets of 600-200 pixels (0.5% and 0.2%, Figure S6). This is independent of the size of the original data, making this method particularly well suited for datasets with large sample numbers (pixels).

The analysis of these images shows that with larger datasets, smaller subsets can be used, and the same quality of embedding can be achieved. The sagittal brain dataset with 123,557 pixels shows similar embedding with 618 pixels in the subset (0.5% subsampling) and 12,356 pixels (10% subsampling) whereas the quality of the segmentation in the coronal and transverse brain images declines rapidly below 1478 and 930 pixels (5 and 10%) respectively. Since the performance is more dependent on the absolute subset size rather than percentage, a subset of at least 2,000 pixels is recommended. Notably, larger subsets are not required for the data containing 1,000,000 pixels compared to 10,000. It is also worth noting that the time taken to perform t-SNE increases quadratically with increasing number of pixels whereas the neural network training is linear, and generally requires much less time (figure S8). Therefore, the rate limiting step in this method remains the application of t-SNE to the subset of data.

The other consideration is how accurately the reconstruction of a spectrum from the t-SNE space can be achieved. In order to evaluate the success of these results single pixel spectra were compared to the returned spectra from their respective RGB points in the NN t-SNE (Figure S9). This was demonstrated on six spectra were randomly selected from the colorectal cancer dataset in Figure 4. In all cases a very high correlation (r^2^ > 0.9) between the original and returned spectra were observed.

The final consideration for this approach is the neural network training parameters. There are a large number of possible combinations that could be altered to varying effect. To exhaustively study this landscape is beyond the scope of this investigation but would be valuable further work, however a preliminary evaluation of the different neural network training algorithms was carried out. Twelve different algorithms were investigated, and the correlation between original t-SNE and NN trained t-SNE was used to evaluate this (Figure S10). It was seen that Bayesian regularisation and Levenberg-Marquardt outperform all other algorithms (r^2^ >0.9 compared to <0.75 for all other methods) with Bayesian regularisation having slightly better performance of the two. This is likely because this method is robust towards overfitting which can occur due to the high dimensionality of MSI and other data^54^.

#### Extension to other dimensionality reduction methods

Finally, this approach is not limited to using t-SNE as the basis for dimensionality reduction. Since the UMAP algorithm has shown recent popularity in many different areas we demonstrate the use of neural network learning of UMAP embedding on the vast chromosome conformation capture data shown in Figure 3, and the mass spectrometry imaging dataset used to evaluate performance shown in Figure S5. As with t-SNE, this deep learning training approach is able to reduce the Hi-C data into 3D showing the same segmentation based primarily intra-chromosome interaction (Figure S11a), with additional hierarchical interactions, and transcriptional activity clustering (Figure S11b and 11c). This approach is generalisable to different dimensionality reduction and demonstrates training of deep learning on non-linear dimensionality reduction. Since the field of non-linear dimensionality reduction is continually growing this approach is applicable to other future developments in this area.

## Conclusions

The NN t-SNE method described overcomes the major limitations of conventional t-SNE. The proposed method can be performed on large datasets, return consistent embedding with the ability to incorporate of new data into the embedded space, and return the driving spectral contributors to the changes observed. This presents a new way to more effectively mine high dimensional data that has large sample numbers. This algorithm can also be applied to the other data types described in Table 1, many of which could benefit from this improvement. In many of the fields where t-SNE is currently applied, such as genomics and proteomics, data is expected to soon exceed the current capacity of existing algorithms. For example, the UK Biobank currently contains genomics data from 487,401 samples, again far greater than conventional t-SNE can analyse^55^. These large-scale initiatives will continue to acquire large cohort datasets from many different techniques, and algorithms need to develop to allow processing and mining of these data. This methodology can also be applied to other dimensionality algorithms such as UMAP. Future considerations for this work include additional comparison of subset sampling methods (such as Sobol sampling) and applications to other areas of research such as large-scale genomics initiatives. We have benchmarked the performance of this approach against using standard t-SNE using the correlation metric between the original t-SNE and NN trained t-SNE with r^2^ > 0.95 between the two approaches both for the forward and reverse transformations.

## Author contributions

A.D. and J.B. designed the scope of research. A.D developed the method. S.A.T. optimised the autoencoders and neural networks and performed 3D registration of the glioblastoma data. R.T.S, K.R, C.N. and A.J.T. acquired MSI data from mouse samples and assisted with interpretation of MSI results. E.E. collected the REIMS data and provided interpretation of these results. A.D.C., A.K.N. and O.J.S developed the colon GEMMs and provided interpretation of these results. Y.P. and M.Y. developed the breast GEMMs and provided interpretation of these results. T.M. supported interpretation of the colon GEMM results. B.Y. supported interpretation of the colon and GEMM breast data. P.G. provided interpretation and additional analysis of the Hi-C data. R.J.G., G.H. and J.S. provided the glioblastoma MSI data. I.S.G., M.Y, R.J.G, S.B, O.J.S, and Z.T. are investigators in the CRUK project and supported the research. J.B. is the principal investigator of the Rosetta Grand Challenge team and led the research. A.D. and J.B. wrote the paper. All authors read the manuscript and provided critical comments.

## Data and materials availability

All data, and code relating to this article will be made available in a public repository.

