## Supplementary materials for "Training a neural network to learn other dimensionality reduction removes data size restrictions in bioinformatics and provides a new route to exploring data representations"

**This PDF file includes:**

Materials and Methods

Figures. S1 to S11

Tables S1 to S2

Materials and Methods

### Data processing

Data processing was performed on an Intel Xeon quad core CPU E5-2680 v3 (2.50 GHz) x2 with 128 GB of RAM and for the subset comparison, on an Intel Xeon CPU E5-2698 v4 (2.2 GHz) x2 with 1 TB of RAM. Data were converted from the proprietary Waters .RAW format into imzML using ProteoWizard(*55*) and imzML converter(*56*), and imported into Matlab (version 2017a and statistics and image processing toolbox, The Math-Works, Inc., Natick, MA, USA) using SpectralAnalysis(*57*). Ion images were generated by integrating intensities across each peak. Mean spectra were generated after preprocessing with interpolated rebinning using a bin width of 0.002 Da(*57*). Once imported into Matlab data subsets were taken by selecting every *n^th^* pixel where *n* is specified by the user (based on memory and time constraints). t-SNE was performed with the Matlab function “tsne” (Statistics toolbox) using the exact algorithm and correlation distance metric(*58*).

Neural network training was performed using the Matlab neural network training tool (Neural network toolbox), using 70% training data, 15% testing data, and 15% validation data, with 1000 max epochs, and Bayesian regularisation as the training method unless otherwise stated. UMAP was performed using the code provided by Meehan *et al. (*<https://www.mathworks.com/matlabcentral/fileexchange/71902>, accessed 25/02/2020) using the default parameters apart from correlation distance metric and reduction to three dimensions.

### Hi-C data

Combined inter- and intrachromosomal contact matrices of human GM12878 cell line generated by (reference https://www.cell.com/fulltext/S0092-8674(14)01497-4) were downloaded from the Gene Expression Omnibus (GEO) using the GSE63525 identifier. The contact matrices used in this study were of 1kb resolution, un-normalised (.RAWobserved files), reads mapping to the genome with MAPQ score > 0. The mitochondrial and sex determination chromosomes (X and Y) were not used in this analysis. Data were loaded in sequentially additively binned into 50kb columns (1kb rows were still retained), and any fully zero rows or columns were removed. Training data was taken every 1000^th^ sample resulting in a total of 2,882 samples. The A/B subcompartment annotation file for GM12878 cell line was downloaded from the same GEO repository (file GSE63525_GM12878_subcompartments.bed.gz). The H3K27ac ChIP-seq data for the GM12878 cell line were obtained from the ENCODE consortium(*59*). The fold-change over control signal from two biological replicates was used (file ENCFF180LKW.bigWig). Parsing of bigWig files and annotation of genomic elements was performed using R packages rtracklayer 1.48 and GenomicRanges 1.40.

### GEMM colon models

All animal experiments were carried out in accordance with UK Home Office regulations (PPL 70/8646), and subject to ethical review by the animal welfare and ethical review board of the University of Glasgow. The genetic alleles used in this study were *Vil*^CreERT2^ (El-Marjou et al), *Apc*^fl^ (Shibata et al 1997), *Kras*^G12D^ (Jackson et al 2001) and *Pten*^fl^ (Suzuki et al 2001). All experiments were carried out in 8-12 week old mice of a pure, inbred C57BL6/J genetic background. Cre-mediated genetic recombination was induced through a single intraperitoneal injection of tamoxifen (80mgkg^-1^) on one occasion (*Vil*^CreERT2^ *Apc*^fl/fl^ *Kras*^G12D/+^ or *Vil*^CreERT2^ *Apc*^fl/fl^ *Kras*^G12D/+^ *Pten*^fl/fl^), or two consecutive days (*Vil*^CreERT2^, *Vil*^CreERT2^ *Apc*^fl/fl^, *Vil*^CreERT2^ *Apc*^fl/fl^ *Pten*^fl/fl^ or *Vil*^CreERT2^ *Kras*^G12D/+^), with animals sacrificed and tissues harvested at 72 or 96 hours post induction respectively. For subsequent analysis by MSI, whole snap-frozen tissue specimens were embedded in a 12.5 % (w/v) solution of carboxymethylcellulose prior to sectioning. For REIMS analysis, intestinal epithelial cells were extracted from the small intestine or colon of these genetically engineered mouse models as described previously (Marsh 2008), snap frozen and stored in Eppendorf tubes at -80°C until analysis.

### MSI analysis

MALDI MSI analysis of the breast and colorectal cancer and sagittal, and coronal mouse brain samples, were acquired on Waters Synapt G2si Q-ToF instruments (Waters, UK) with a MALDI source for the breast and colorectal cancer samples, and a prototype uMALDI source for the sagittal mouse brain samples. The transverse mouse brain data were acquired on a Waters Xevo G2-XS Q-ToF instrument with a DESI ion source (Waters, UK) and the glioblastoma samples were acquired using a Bruker RapiFlex instrument (Bruker, Bremen, Germany). A full summary of experimental prarmeters for MSI data is provided in table S2.

### REIMS analysis

REIMS of the cell extracts was performed using a Waters Xevo G2-XS operated in negative ion mode, 50 - 1500 *m/z* at 2 scans/s, using irrigated bipolar forceps for sample mobilisation. An Erbe VIO 50C electrosurgical generator was operated in bipolar mode at an output power of 15 W. The aerosol generated from the sample was aspirated and transferred to the REIMS source using Tygon tubing where it was mixed with propan-2-ol in a 't-piece' prior to introduction to the mass spectrometer.

Figure S1. Results of NN t-SNE on Hi-C data labelled by the H3K27ac ChIP-seq signal (epigenetic mark linked to active gene expression). Regions with high transcriptional activity (yellow) are clustered together in the 3D space.

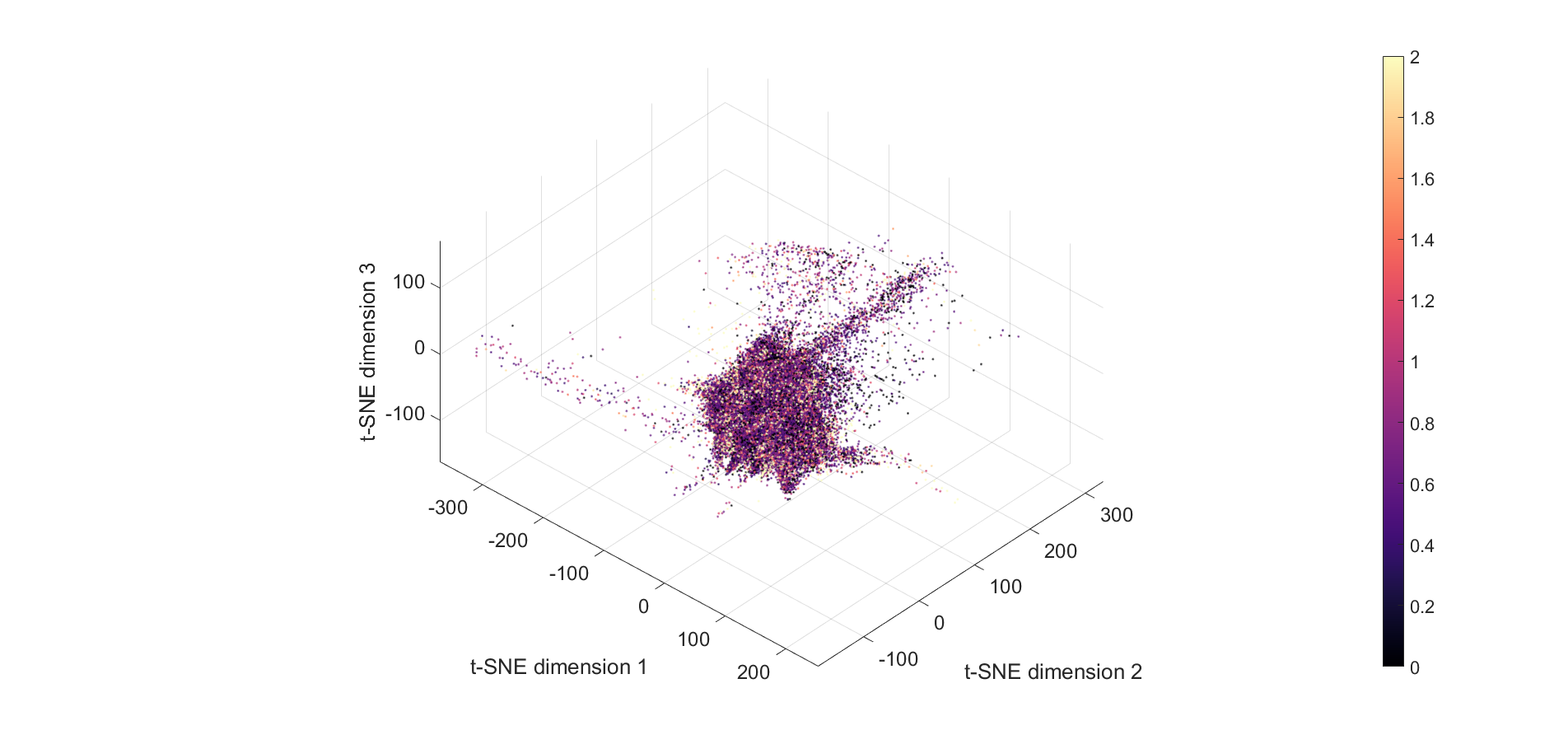

Figure S2. Results of NN-tSNE trained on the 21^st^ section (highlighted in red) of a 3D brain dataset which does not contain glioblastoma (which is later segmented in dark blue) and subsequently applied to the data from the remaining sections showing a consistent segmentation of the similar anatomy. More importantly the pixels from the glioblastoma are identified as being different from any data used in the training process. Sections are ordered by depth left to right and bottom to top, and the approximate corresponding image from the Allen brain atlas is sown below each image.

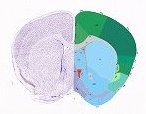

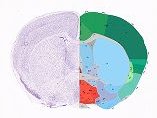

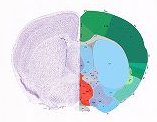

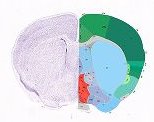

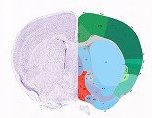

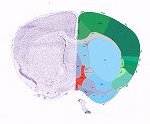

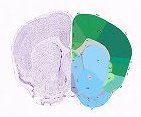

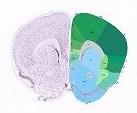

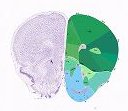

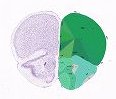

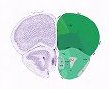

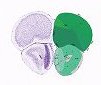

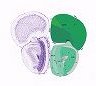

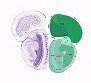

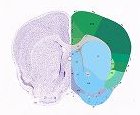

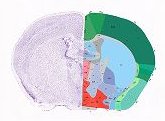

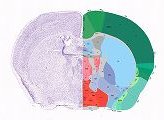

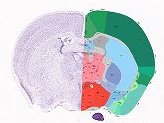

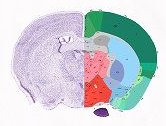

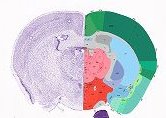

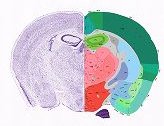

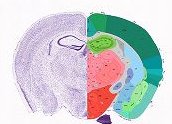

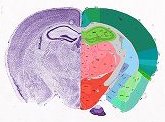

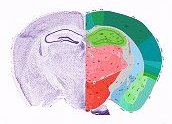

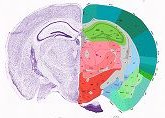

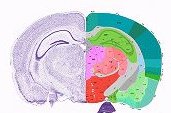

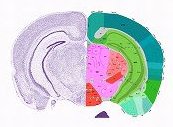

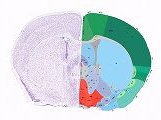

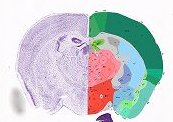

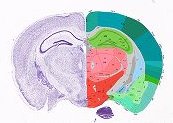

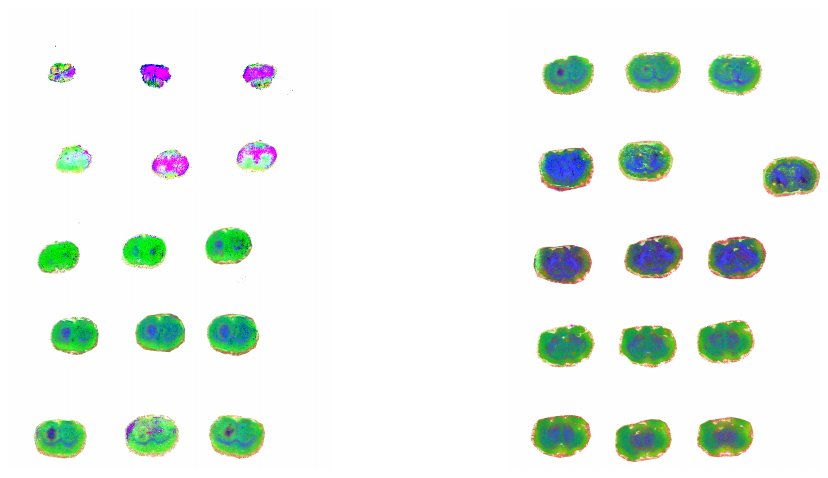

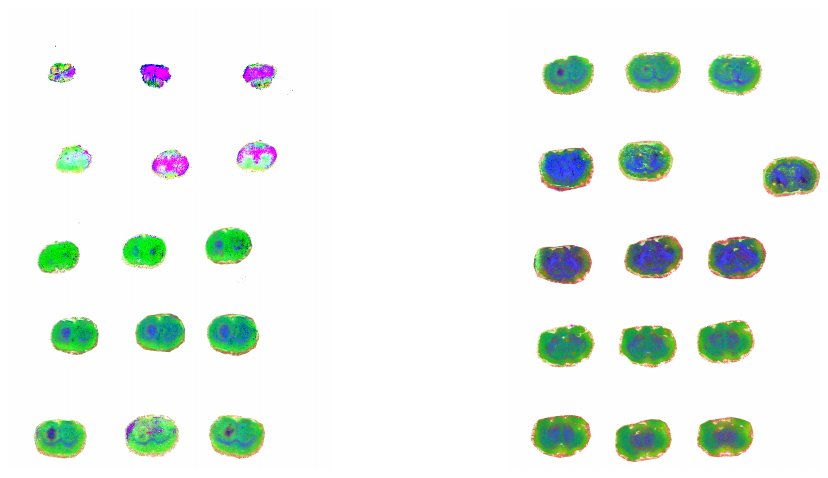

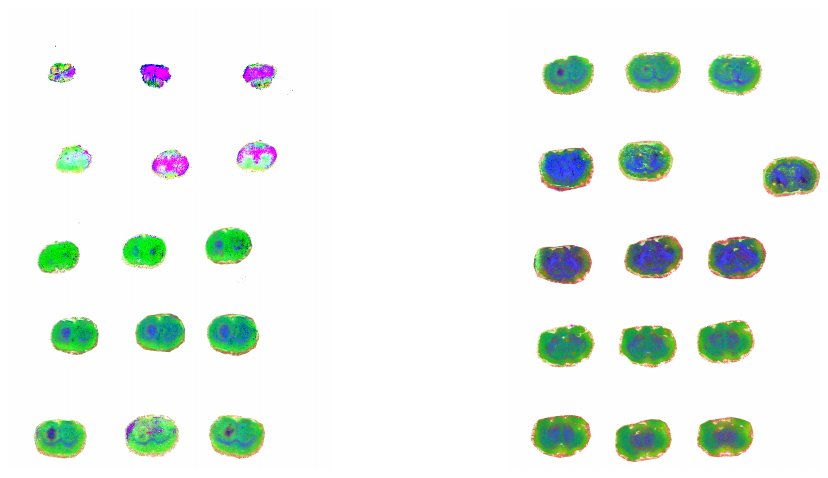

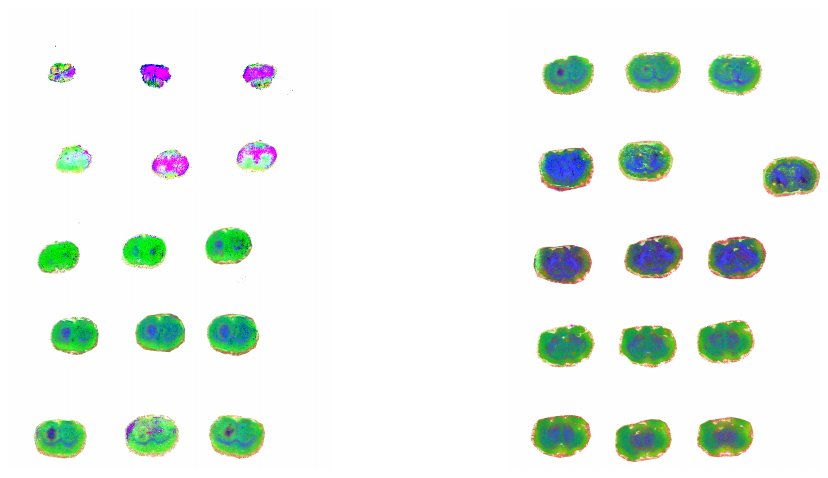

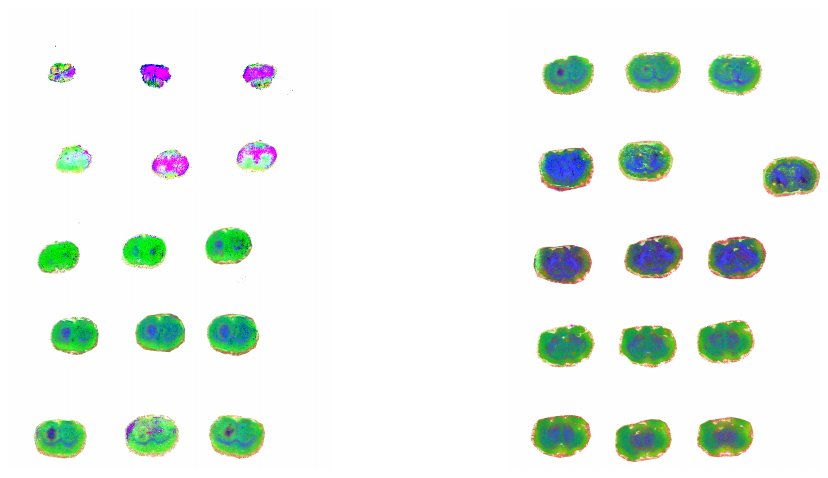

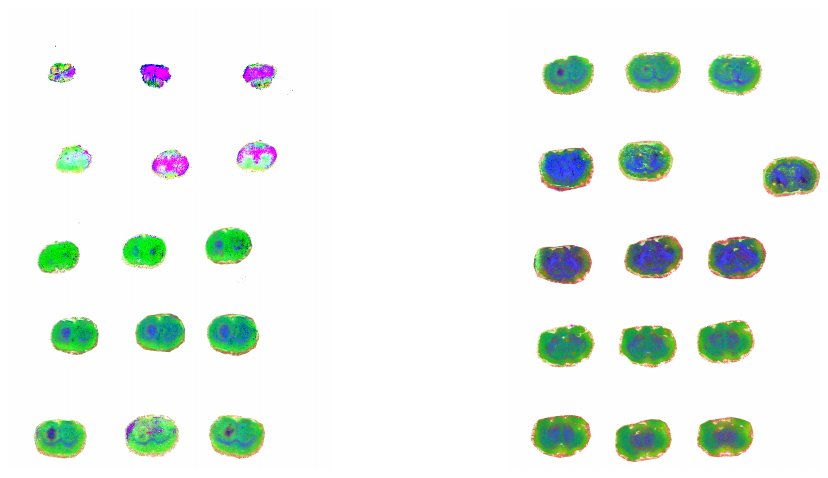

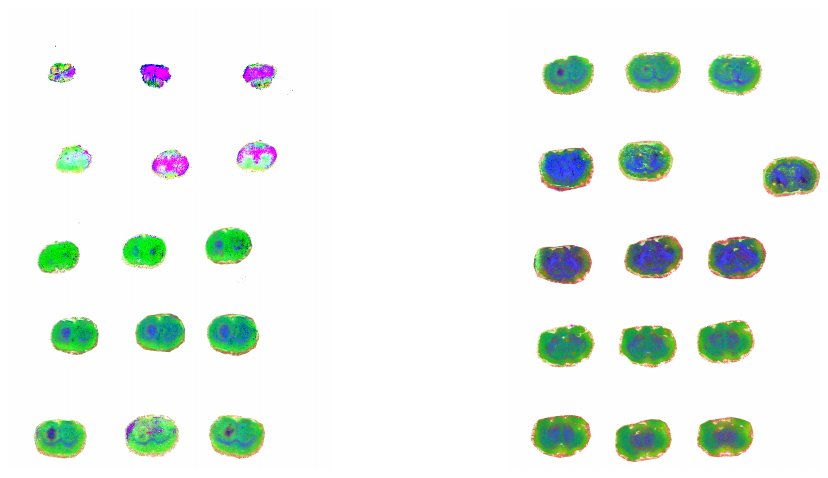

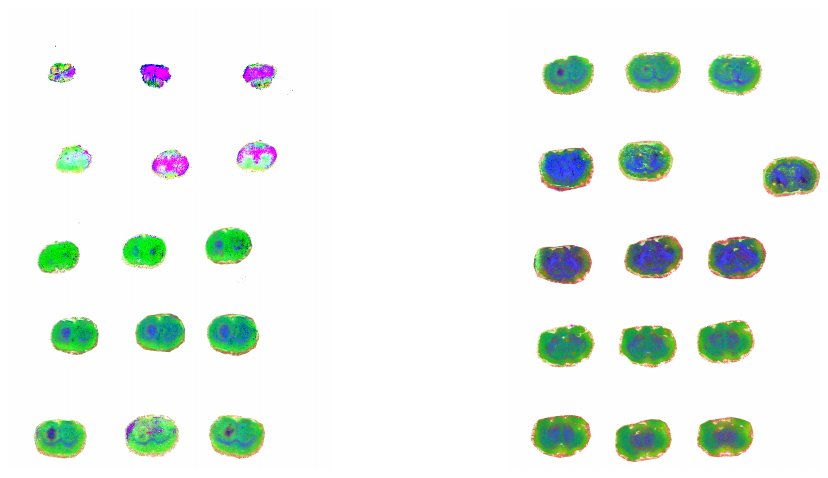

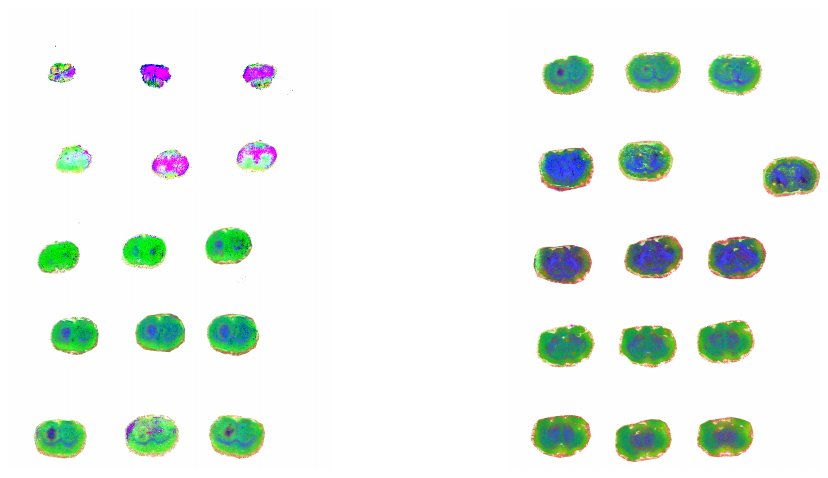

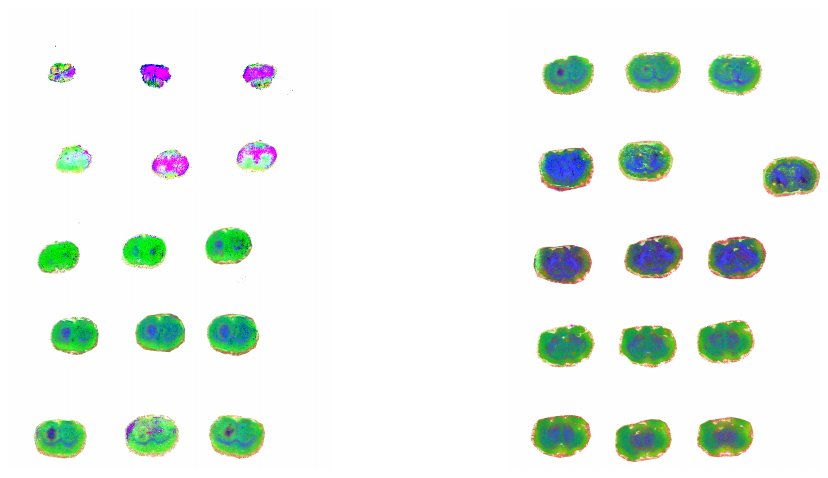

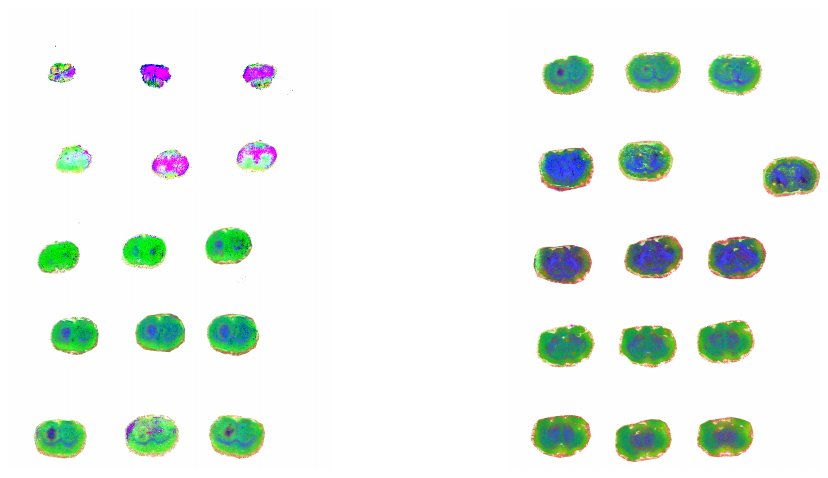

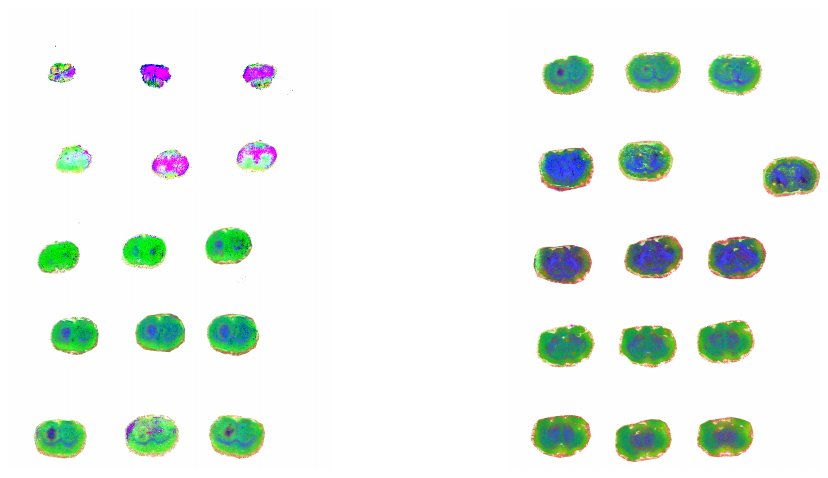

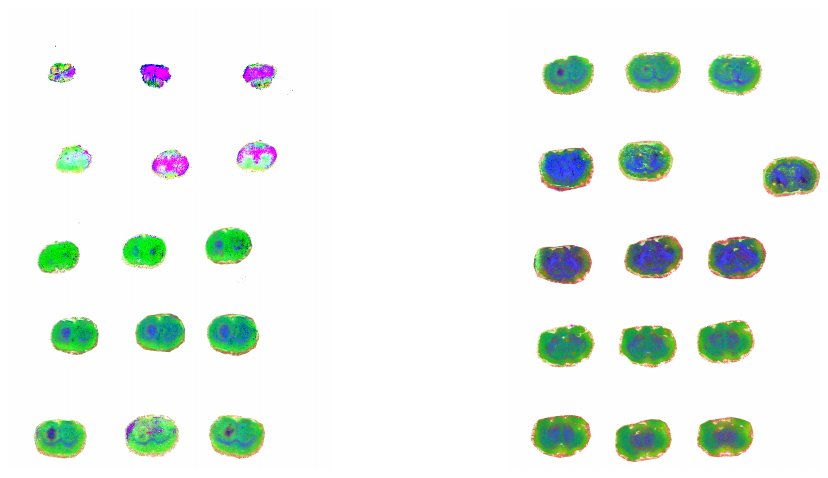

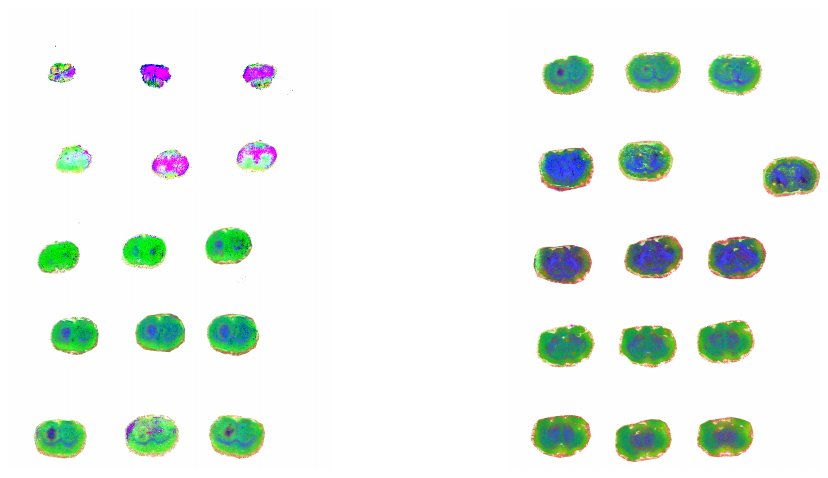

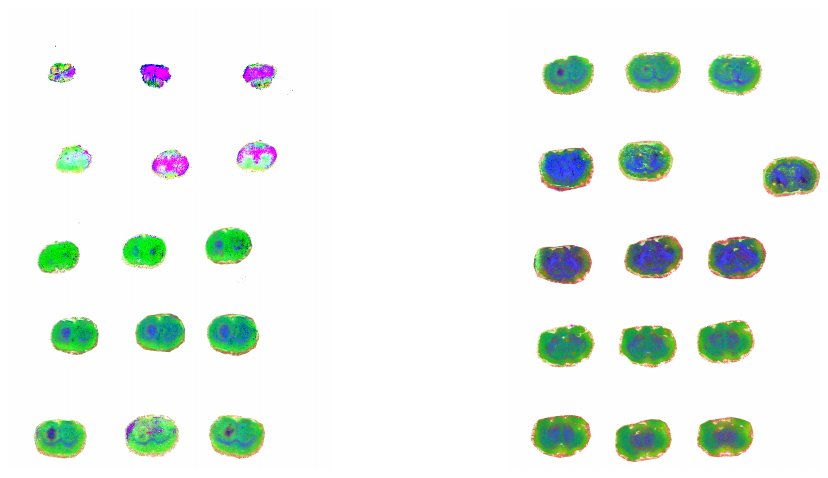

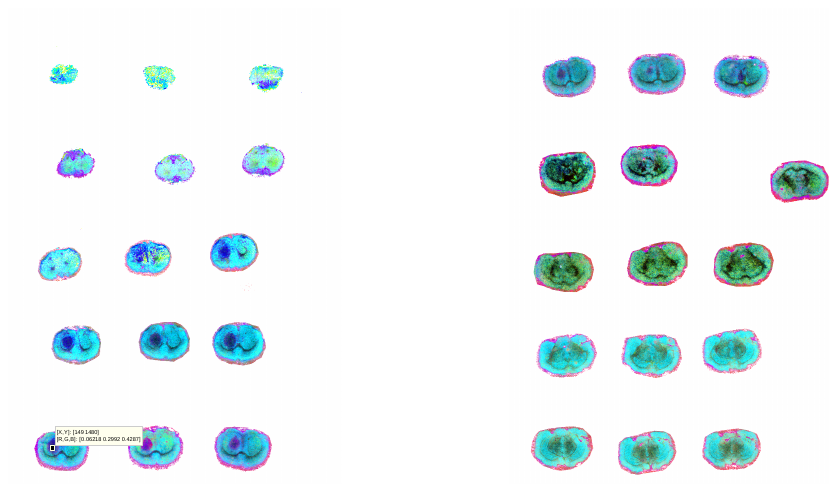

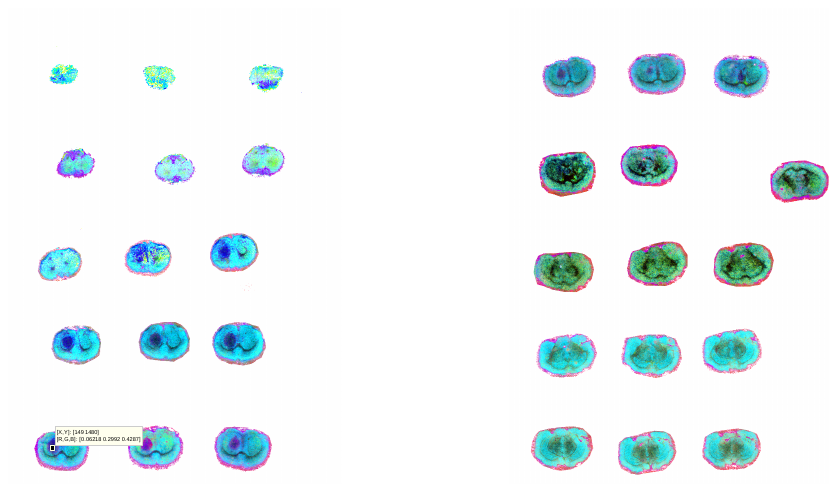

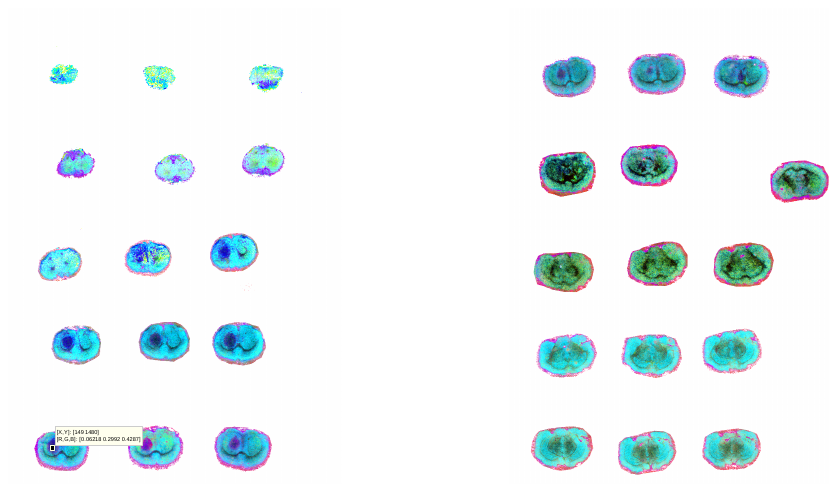

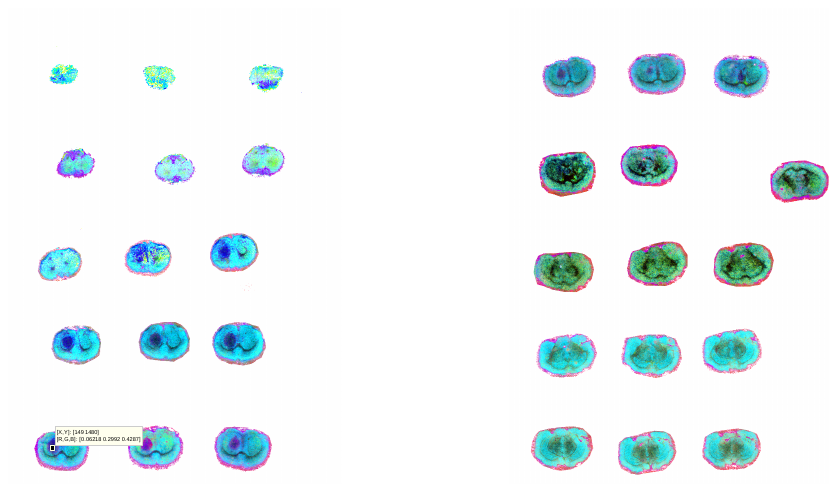

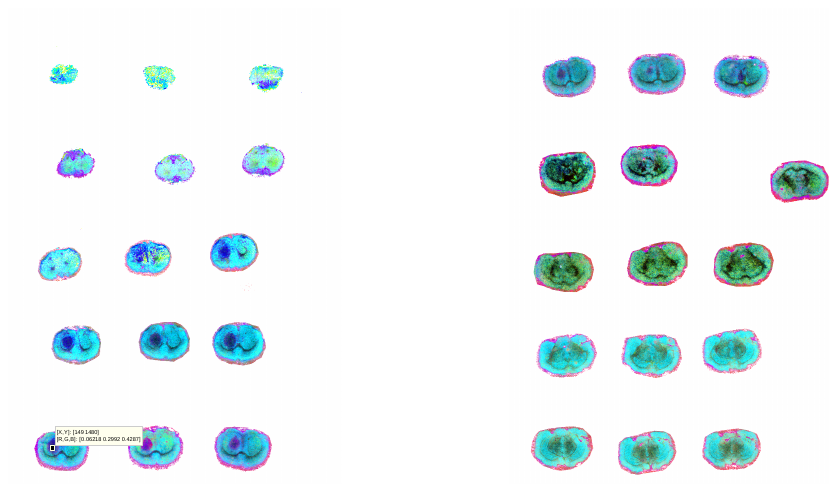

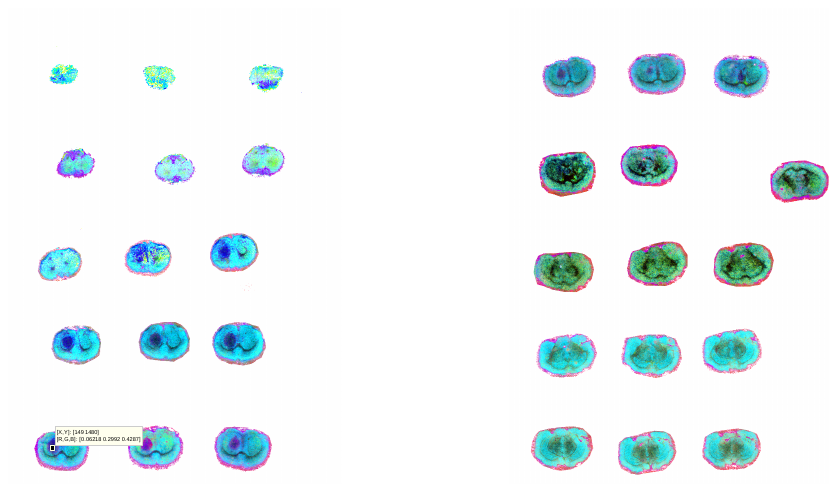

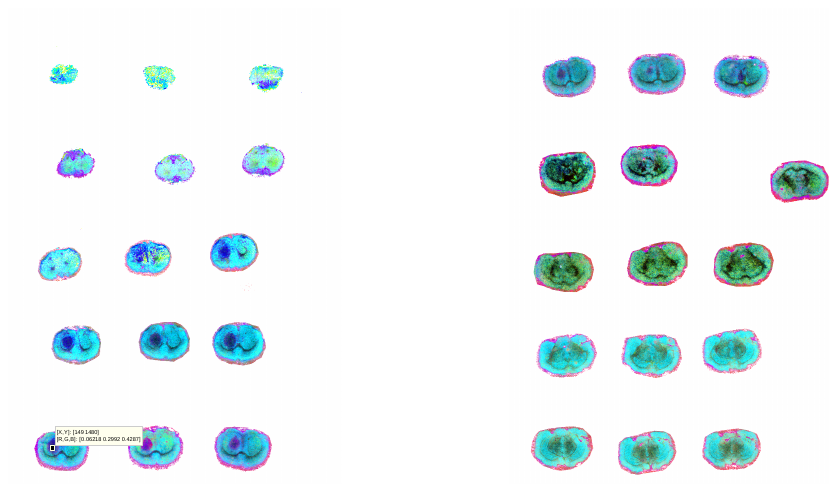

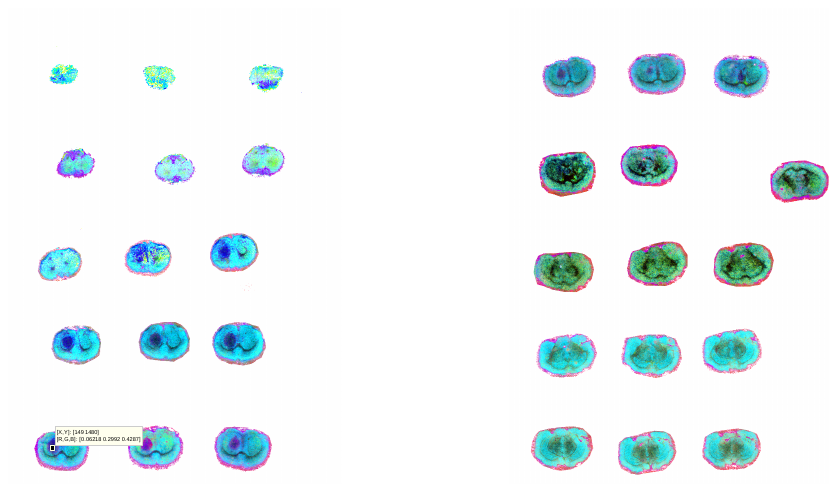

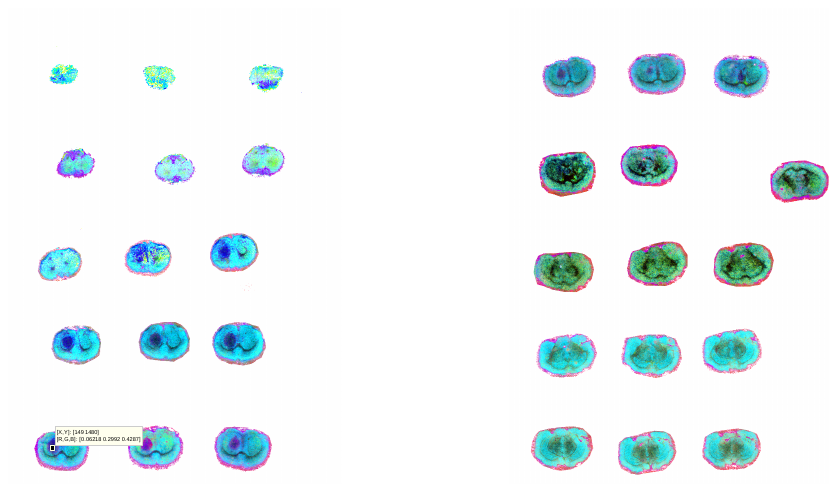

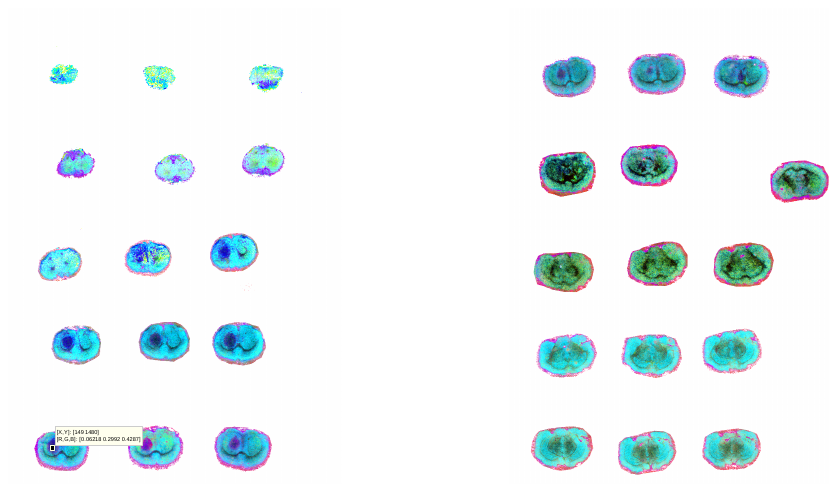

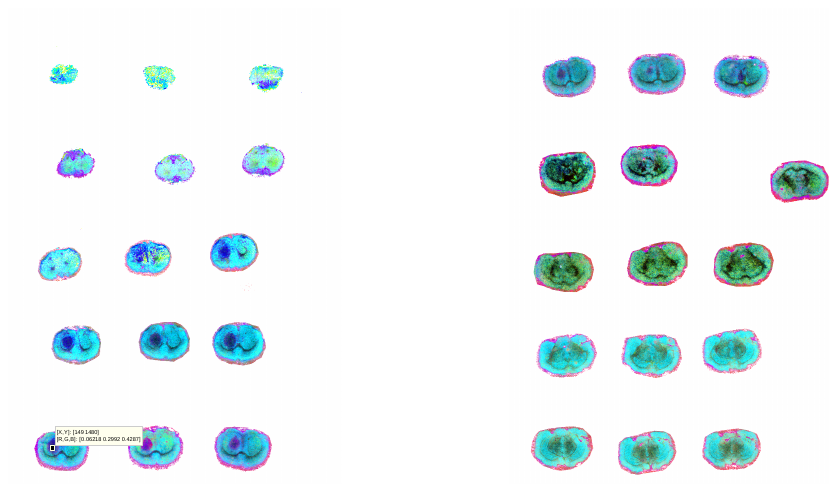

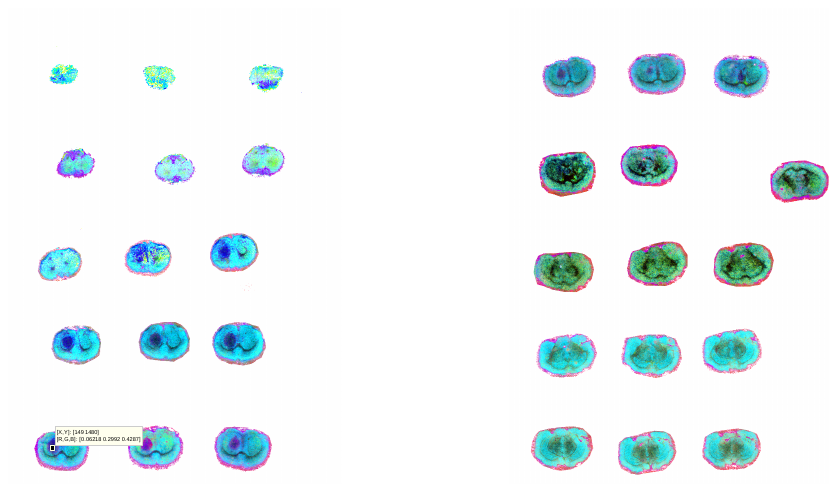

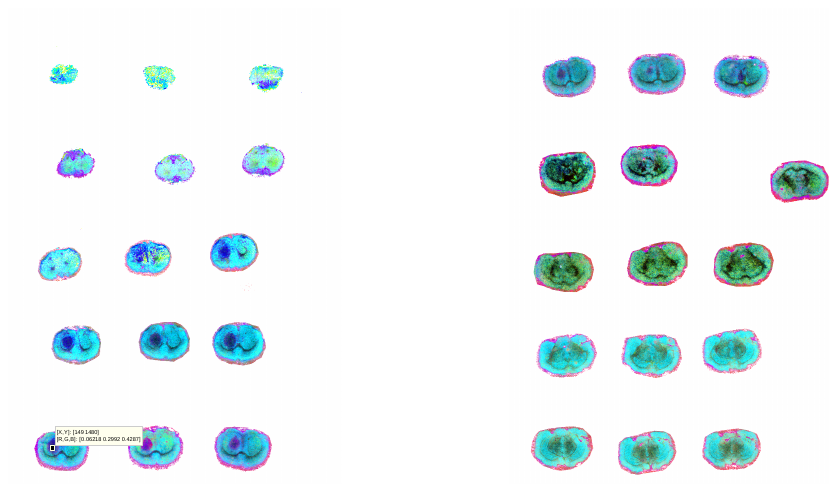

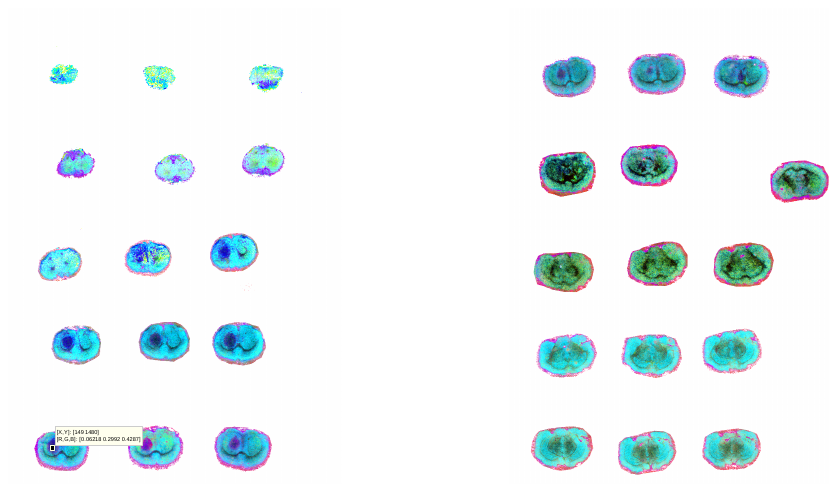

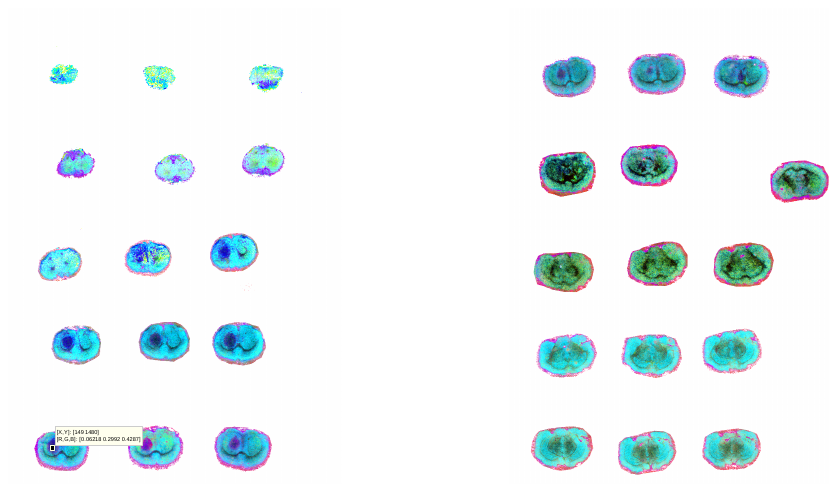

Figure S3. Comparison of NN-tSNE and PCA applied to REIMS data from cell pellets collected from colon (x) and small intestine (o) from genetic mouse models. An additional nine blinded cell pellets were also collected (black) which were used to evaluate the classification using these approaches. NN t-SNE shows much clearer segmentation of the different genotypes based on their metabolic profiles than PCA.

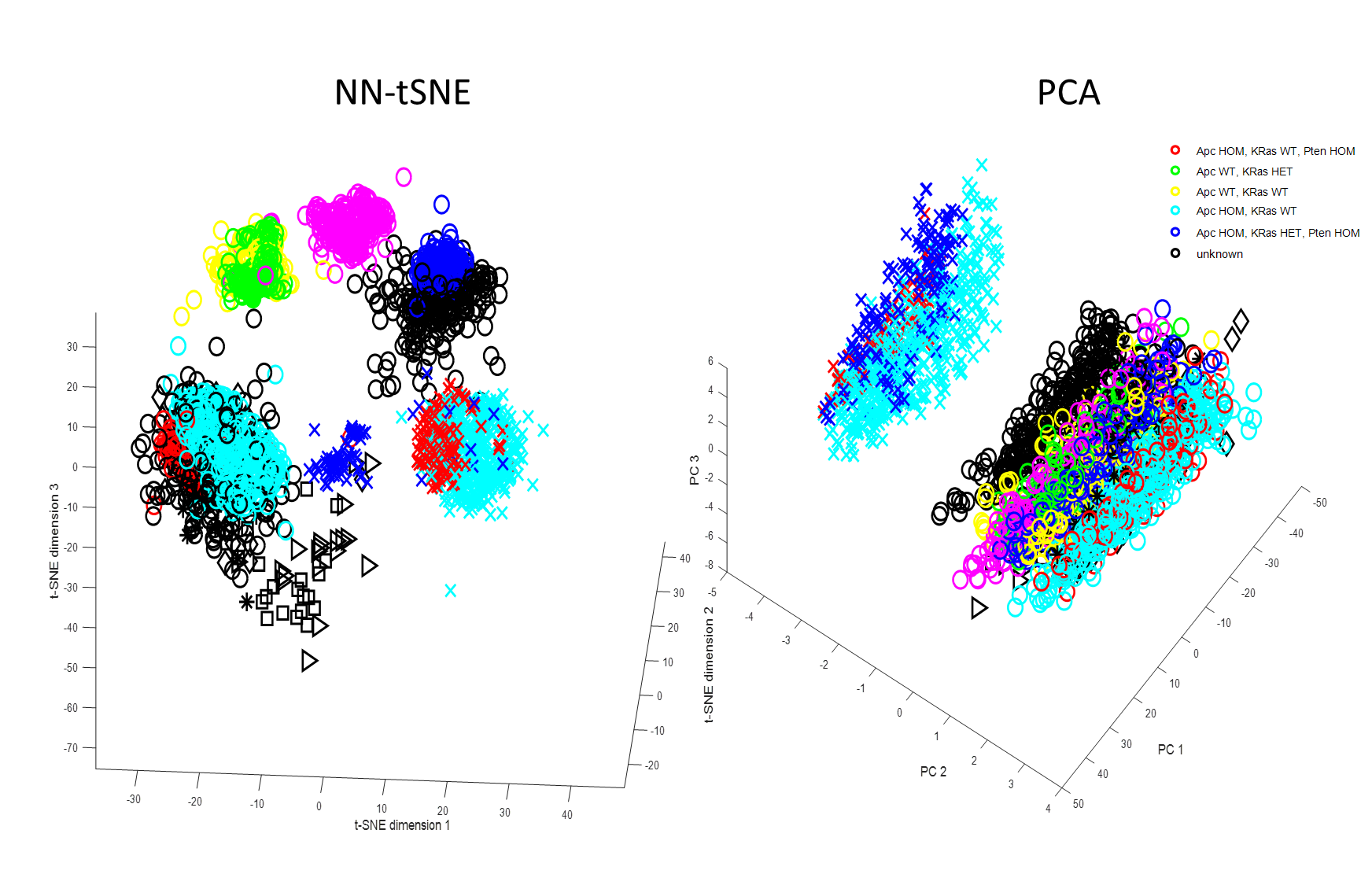

Table S1. Comparison of classification results using either PCA or NN-tSNE as dimensionality reduction prior to LDA classification training. Classification on the NN-tSNE reduced data outperforms the PCA reduced in all metrics on these data.

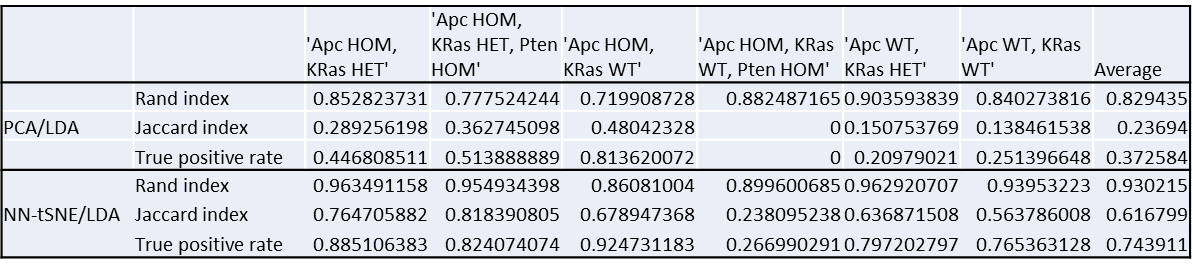

Figure S4. A comparison of ordered vs. random subsampling on the accuracy of the segmentation as evaluated by comparing the correlation to the t-SNE results. This shows no difference in these two sampling methods.

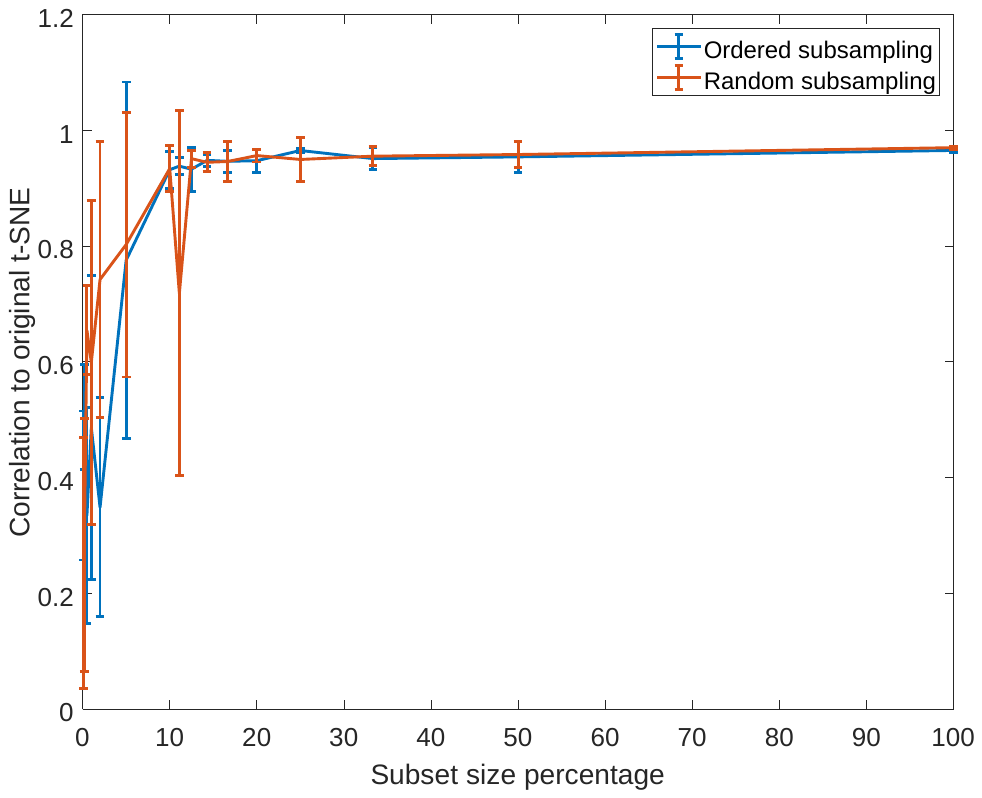

Figure S5. Comparison of the random and ordered subsampling on the resulting images from the sagittal brain dataset. These results show no major differences in the anatomical features obtained using these two different sampling approaches.

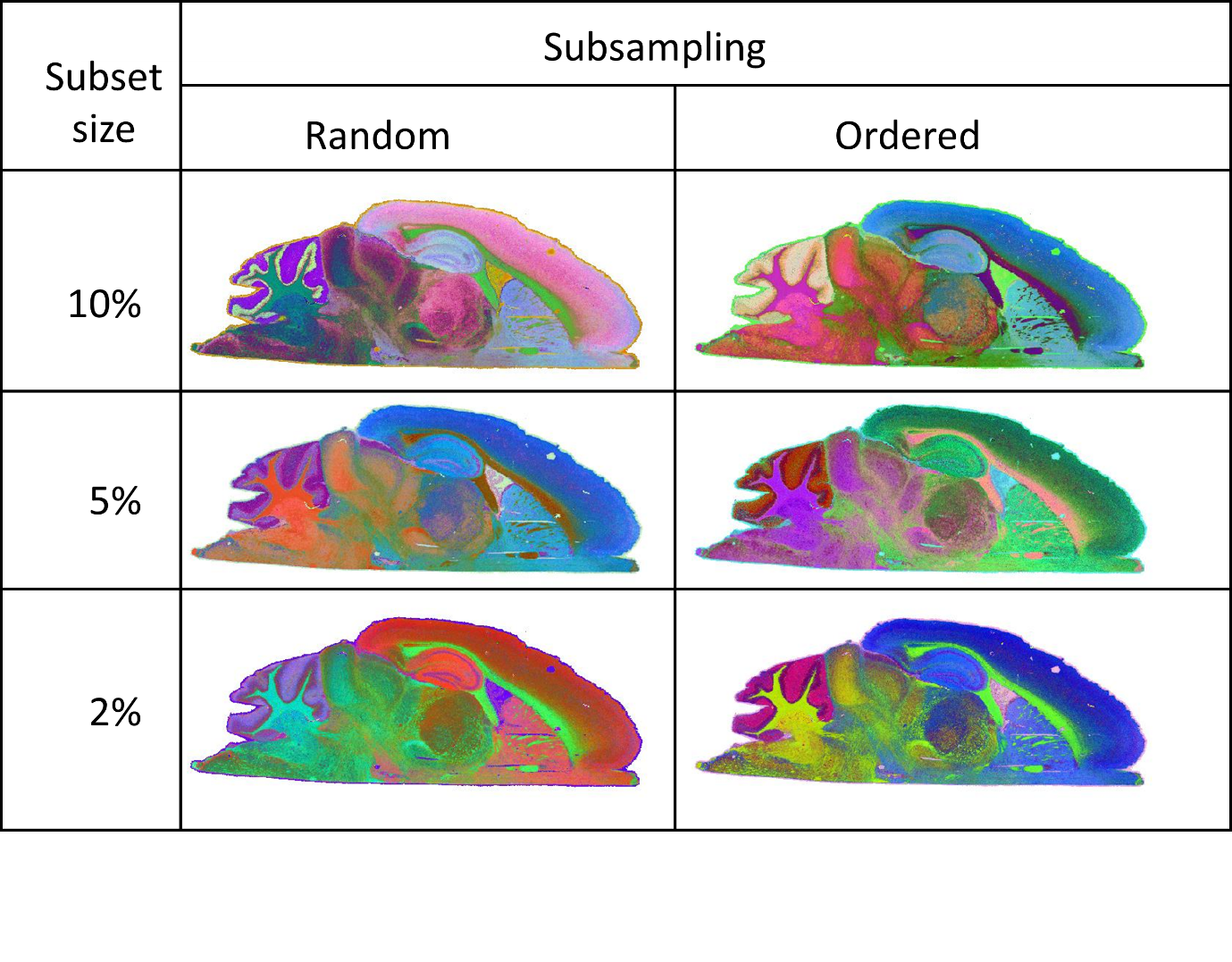

Figure S6. Comparison of the NN-tSNE embedding on the data when t-SNE is performed on different sized subsets. The threshold at which the anatomical feature segmentation is dependent on the number of pixels rather than the percentage of the original data size.

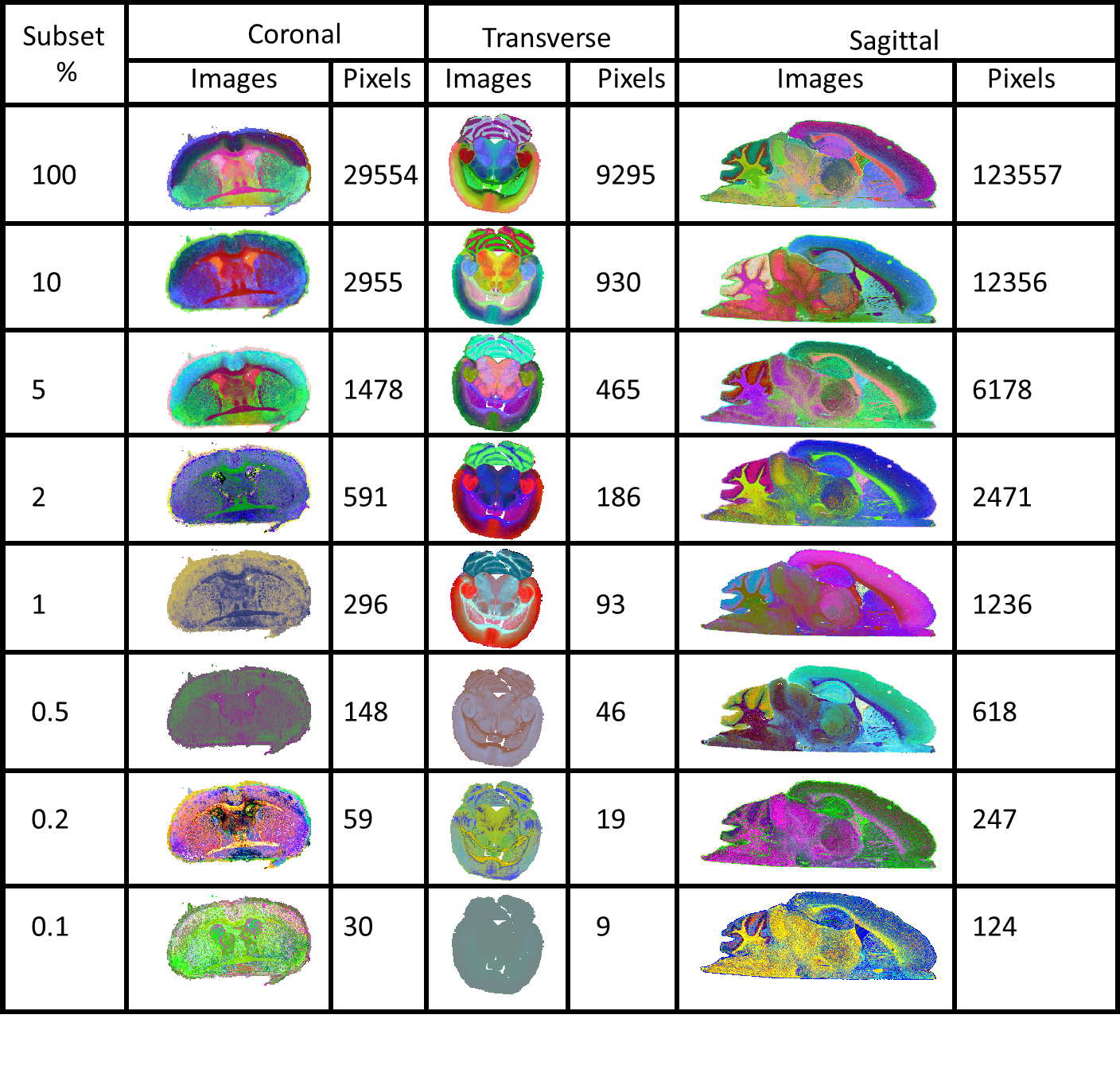

Figure S7. Autocorrelation applied to measure the influence of subset size on the image quality of the NN-tSNE results on the sagittal mouse brain data.

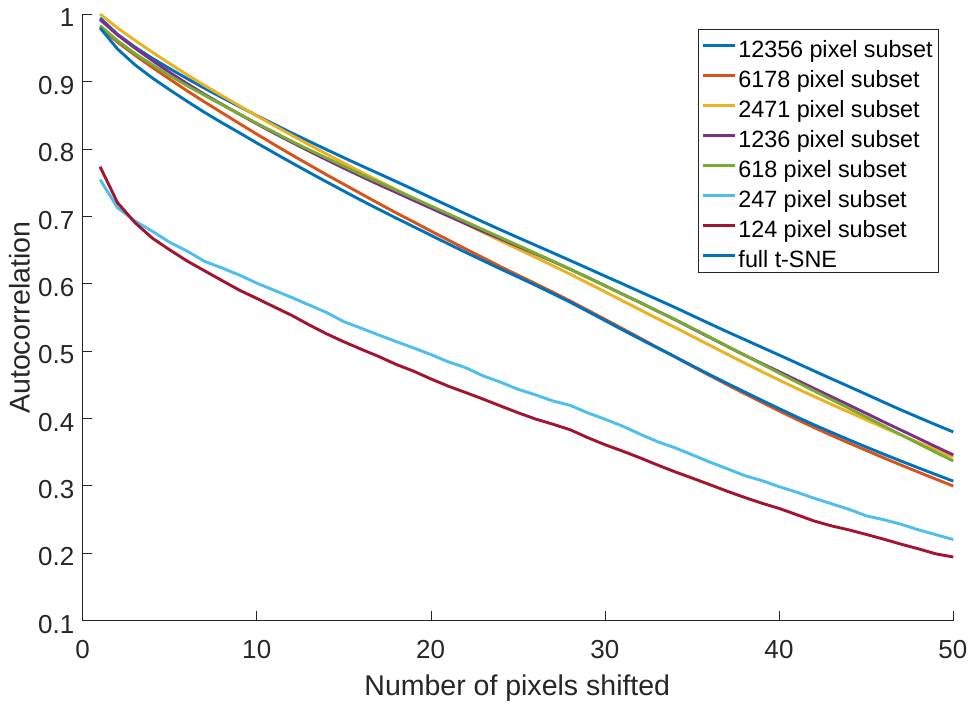

Figure S8. Comparison of the time taken to perform t-SNE and neural network training on different sized datasets. Above 10,000 pixels this becomes prohibitively slow to run as a routine analysis, whereas by preforming NN-tSNE, this can be reduced to a much more feasible timeframe. Of note, to run NN t-SNE on over 10,000 pixels you would still only need to train the data using 2,000 pixels (taking around 3 minutes).

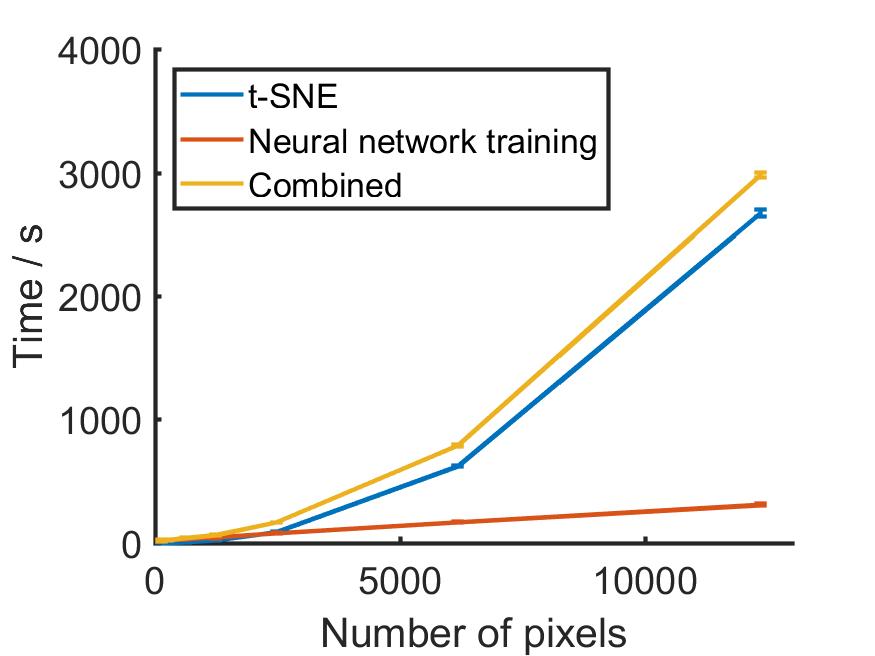

Figure S9. Comparison of the returned spectral contribution from an actual spectrum, as compared to their original spectra. Each one shows a high correlation to the spectrum that it was derived from.

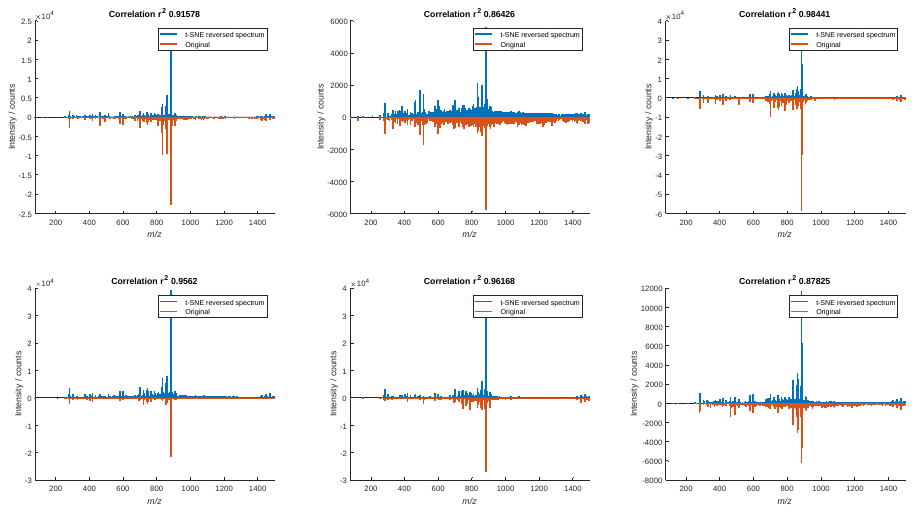

Figure S10. Comparison of the different neural network training methods applied to the MSI image of sagittal mouse brain evaluated on the correlation to the original t-SNE and the anatomical feature identification. The Levenberg-Marquardt and Bayesian regularisation methods are far superior to all other training approaches.

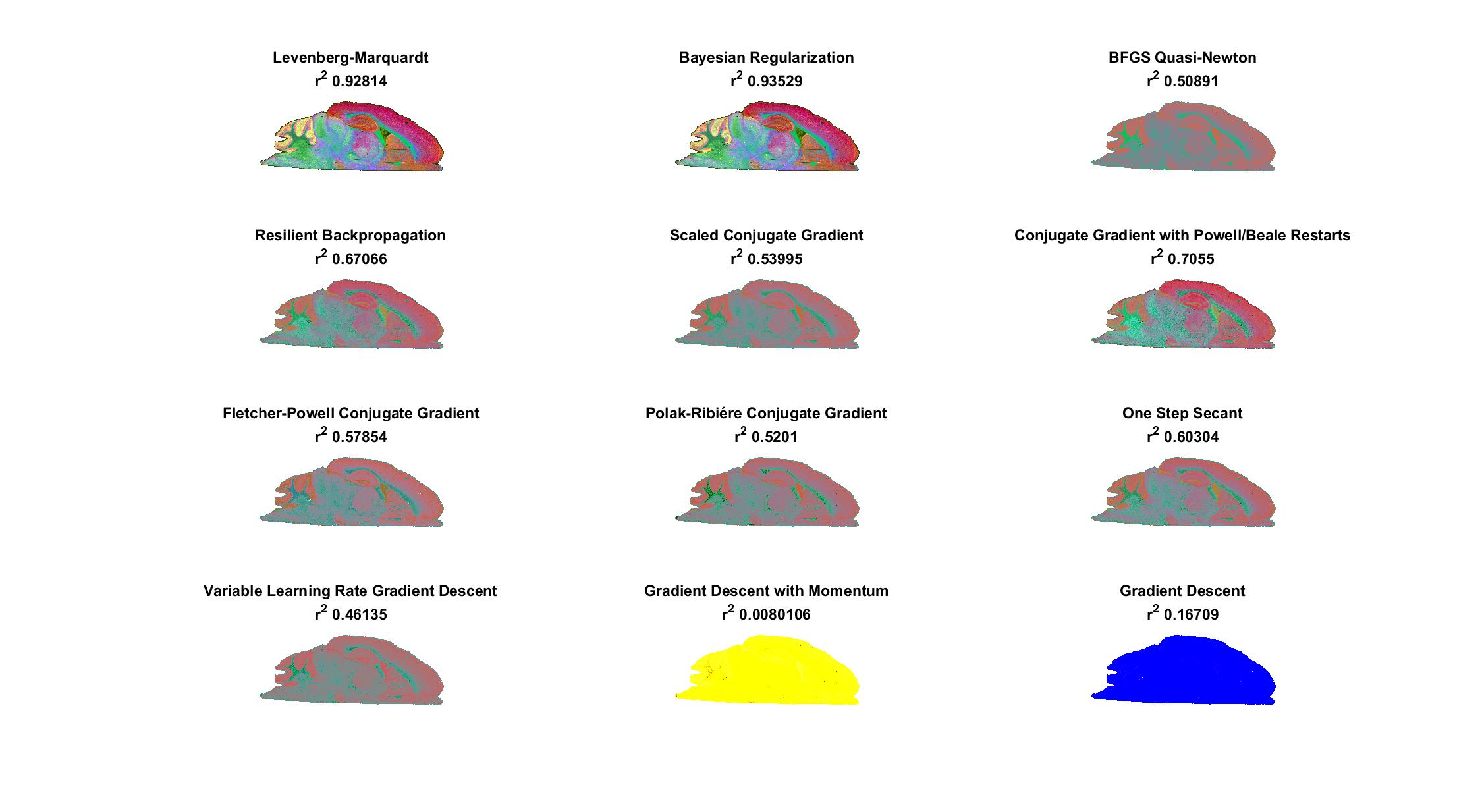

Figure S11. Results of neural network training on UMAP applied to a subset of the Hi-C data seen in figure 3. As with the neural network t-SNE the NN-UMAP primarily segments the data based on chromosome number (a), with other detail being discerned such as the A/B sub-compartments (b) and co-clustering transcriptional activity (c).

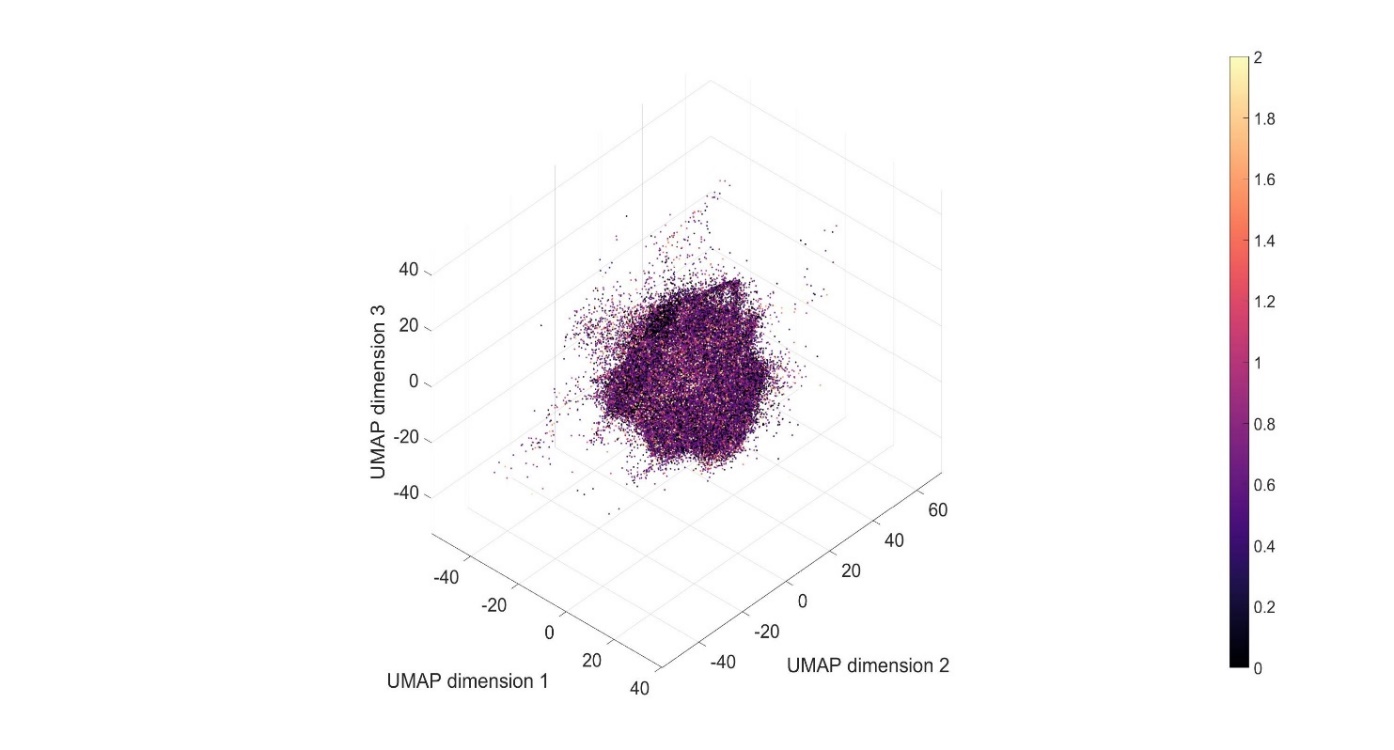

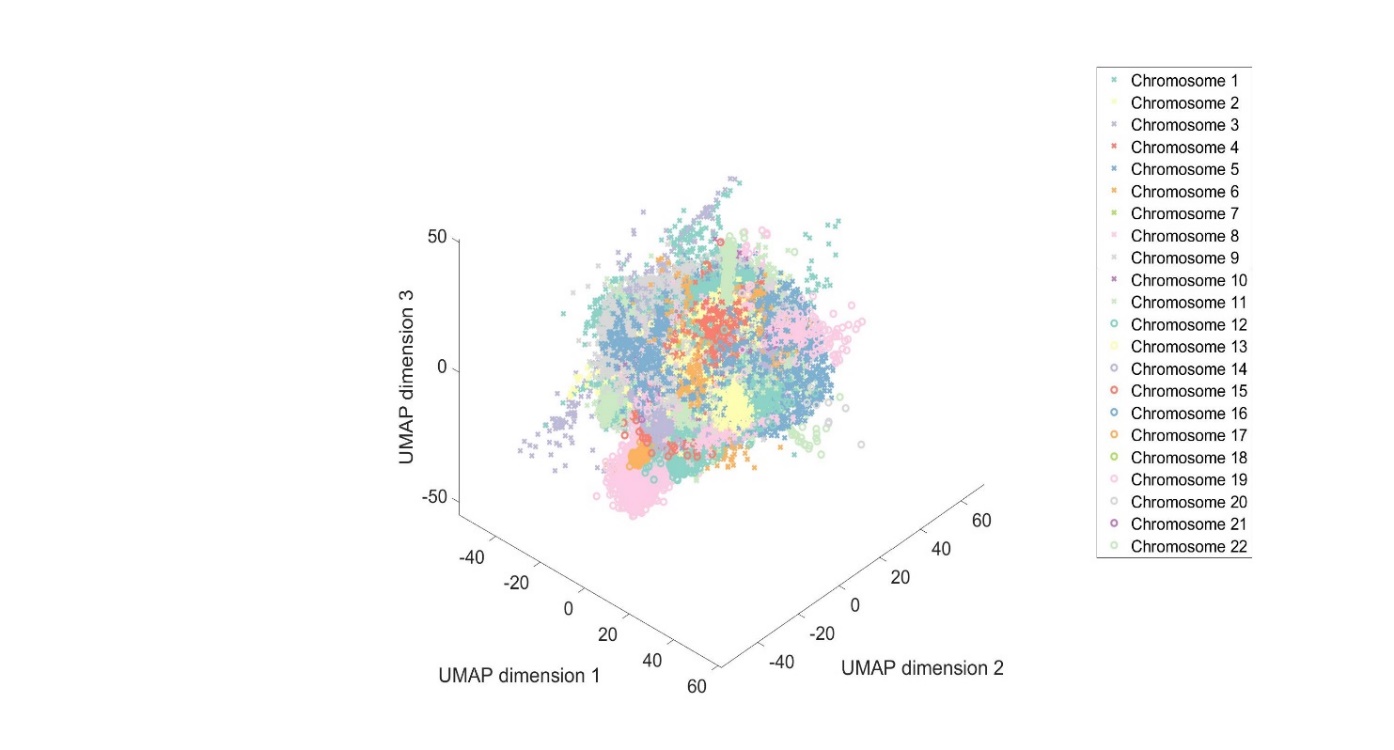

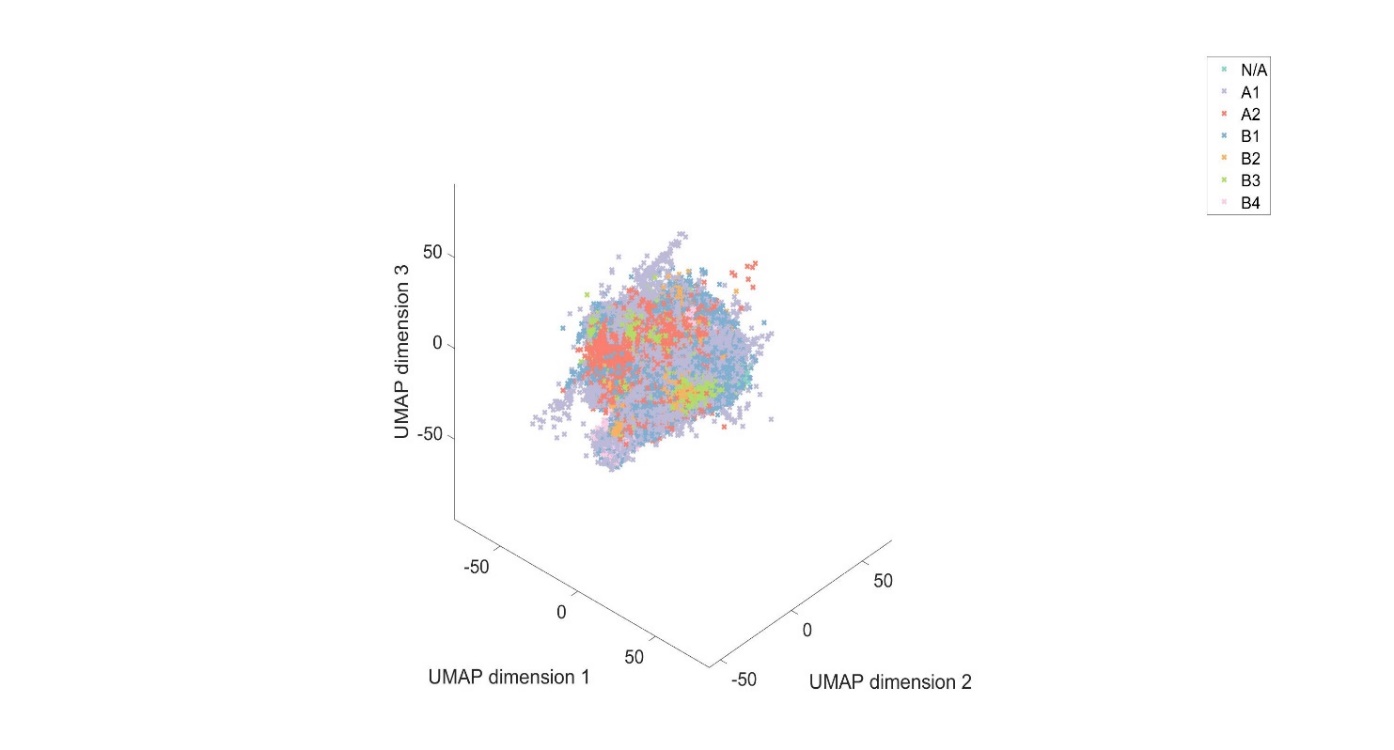

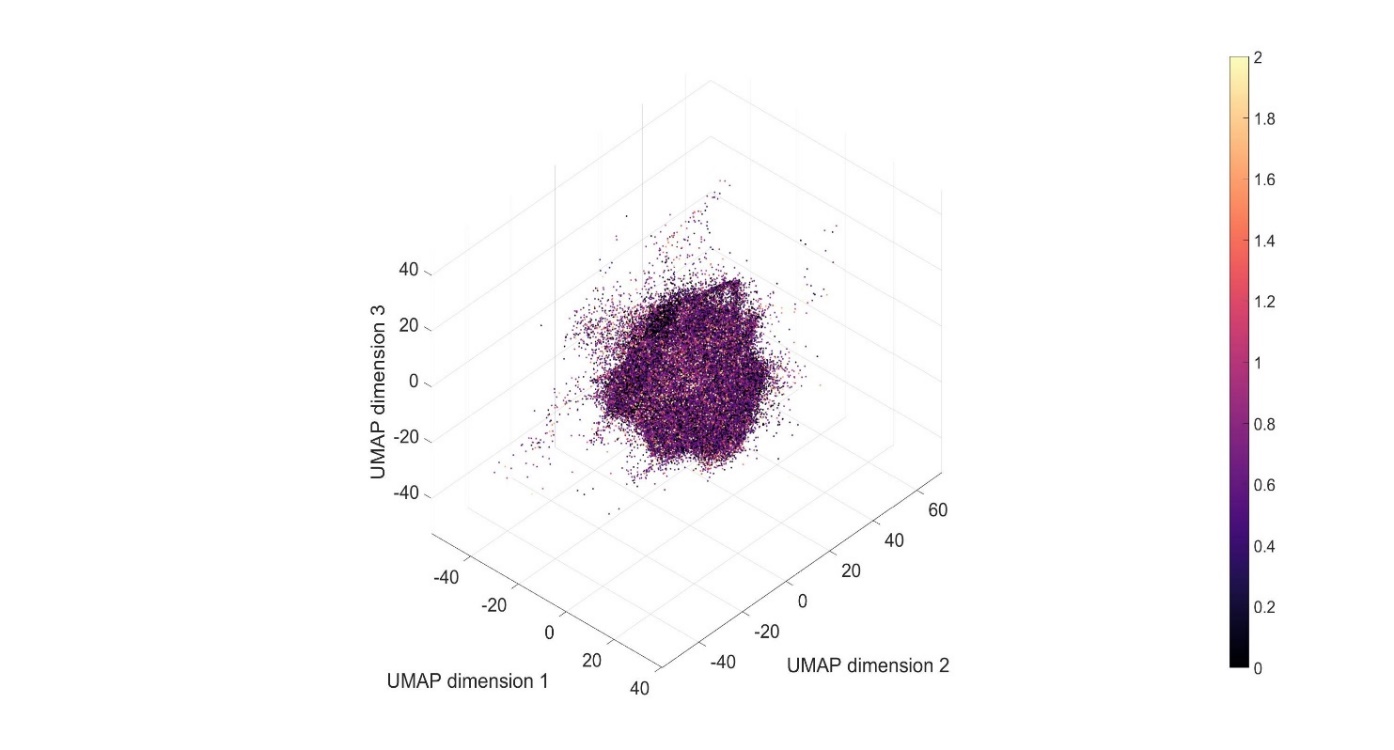

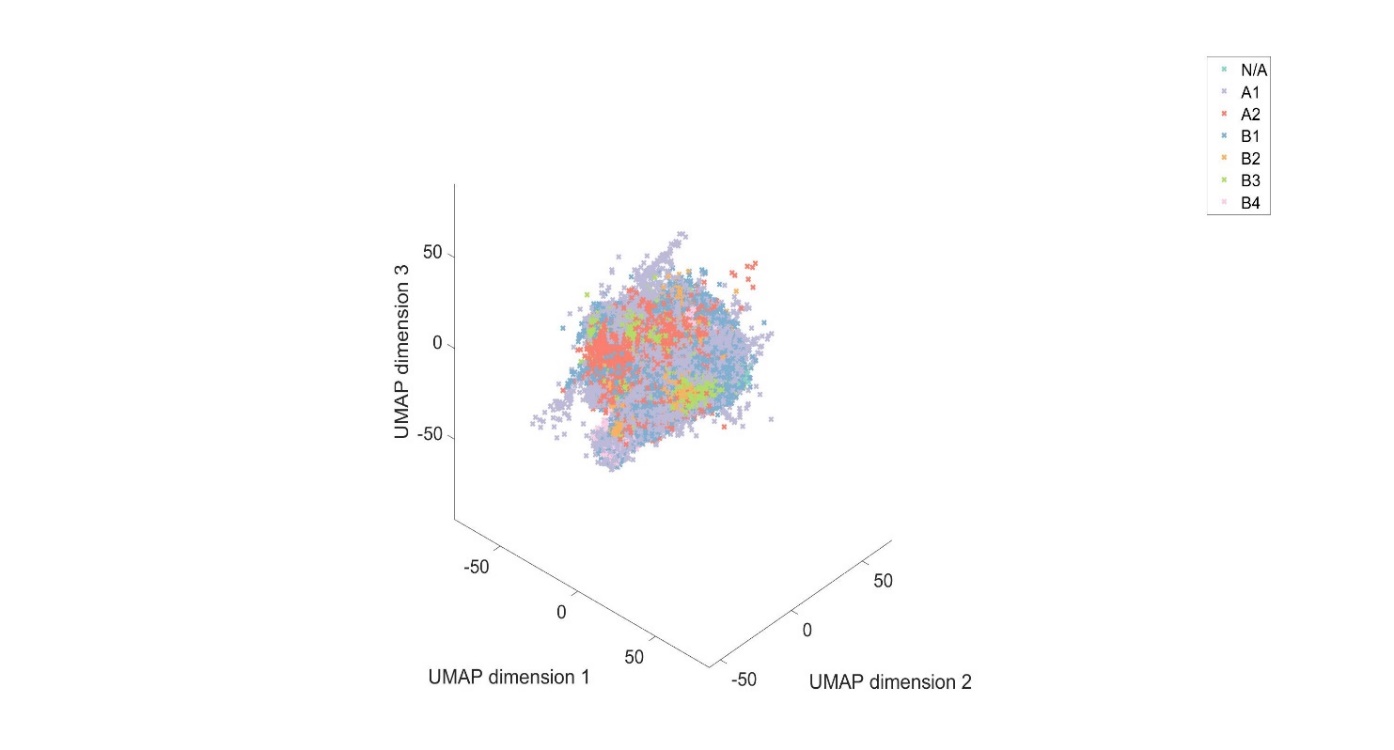

Table S2. Summary of experimental parameters for the acquisition of the mass spectrometry imaging datasets.

| **Dataset** | **Ionisation source** | **Pixel size / μm** | ***m/z* Range** | **Matrix/solvent** | **Scan speed / pixels per second** | **Polarity** |
| --- | --- | --- | --- | --- | --- | --- |
| Breast cancer model | DESI | 75 | 50-1500 | 95/5 MeOH/water 0.02 mg/ml raffinose | 2 | Negative |
| Sagittal brain | MALDI | 20 | 100-1500 | CHCA | 20 | Positive |
| Transverse brain | DESI | 50 | 50-1200 | 95/5 MeOH/water | 4 | Positive |
| Coronal brain | MALDI | 45 | 100-1200 | CHCA | 2 | Positive |
| Colorectal cancer model | MALDI | 50 | 50-1500 | 9-AA | 2.5 | Negative |
| Glioblastoma | MALDI | 50 | 100-1000 | 2,5-DHB | 25 | Positive |
